# Neutralizing IFNL3 Autoantibodies in Severe COVID-19 Identified Using Molecular Indexing of Proteins by Self-Assembly

**DOI:** 10.1101/2021.03.02.432977

**Authors:** Joel J. Credle, Jonathan Gunn, Puwanat Sangkhapreecha, Daniel R. Monaco, Xuwen Alice Zheng, Hung-Ji Tsai, Azaan Wilbon, William R. Morgenlander, Yi Dong, Sahana Jayaraman, Lorenzo Tosi, Biju Parekkadan, Alan N. Baer, Mario Roederer, Evan M. Bloch, Aaron A. R. Tobian, Israel Zyskind, Jonathan I. Silverberg, Avi Z. Rosenberg, Andrea L. Cox, Tom Lloyd, Andrew L. Mammen, H. Benjamin Larman

## Abstract

Unbiased antibody profiling can identify the targets of an immune reaction. A number of likely pathogenic autoreactive antibodies have been associated with life-threatening SARS-CoV-2 infection; yet, many additional autoantibodies likely remain unknown. Here we present Molecular Indexing of Proteins by Self Assembly (MIPSA), a technique that produces ORFeome-scale libraries of proteins covalently coupled to uniquely identifying DNA barcodes for analysis by sequencing. We used MIPSA to profile circulating autoantibodies from 55 patients with severe COVID-19 against 11,076 DNA-barcoded proteins of the human ORFeome library. MIPSA identified previously known autoreactivities, and also detected undescribed neutralizing interferon lambda 3 (IFN-λ3) autoantibodies. At-risk individuals with anti-IFN-λ3 antibodies may benefit from interferon supplementation therapies, such as those currently undergoing clinical evaluation.

**One-Sentence Summary:** Molecular Indexing of Proteins by Self Assembly (MIPSA) identifies neutralizing IFNL3 autoantibodies in patients with severe COVID-19.

**Graphical Abstract:** 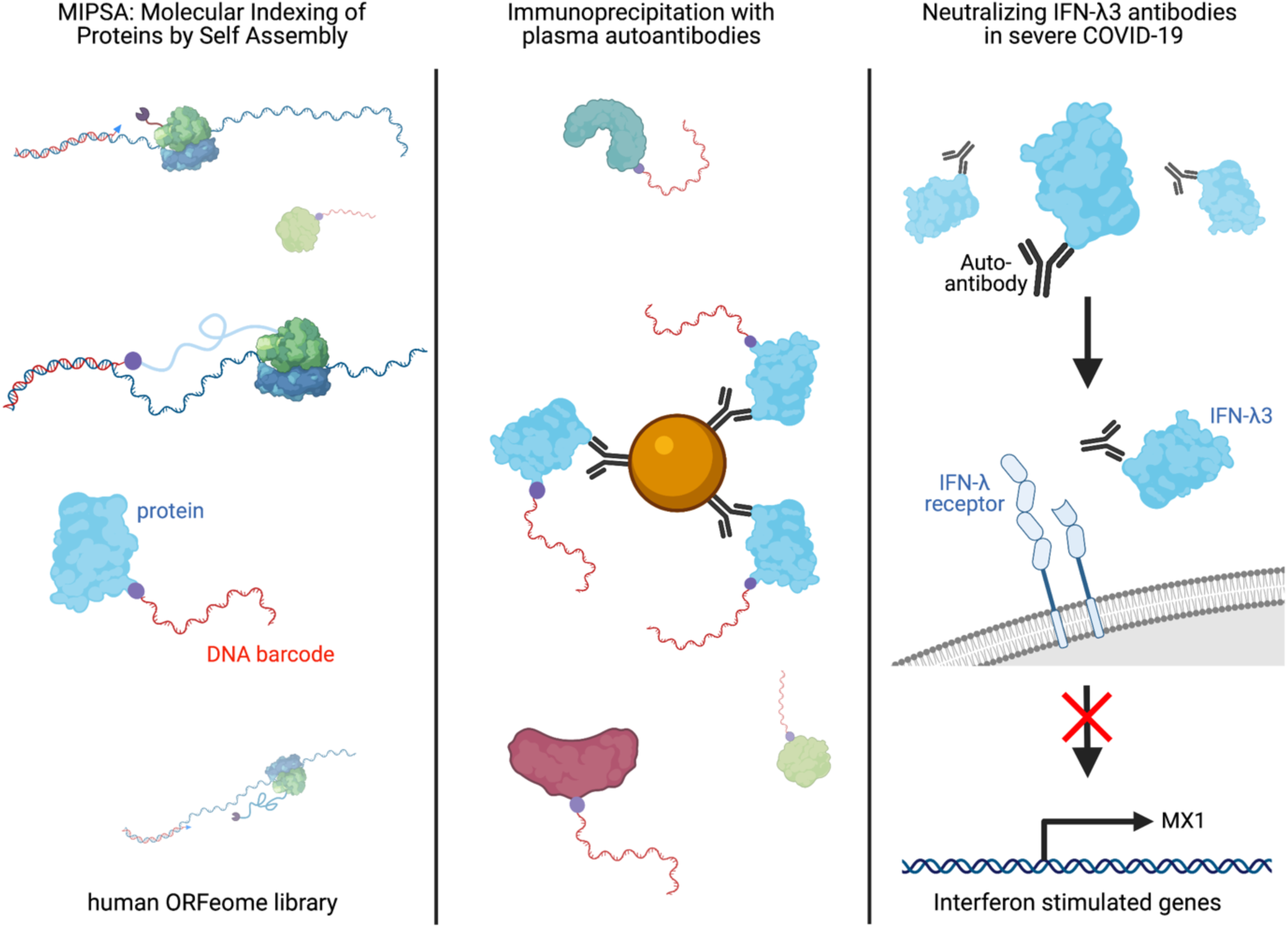

## Introduction

Unbiased analysis of antibody binding specificities can provide important insights into health and disease states. We and others have utilized programmable phage display libraries to identify novel autoantibodies, characterize anti-viral immunity and profile allergen-specific IgE antibodies.(*1–4*) While phage display has been useful for these and many other applications, most protein-protein, protein-antibody and protein-small molecule interactions require a degree of conformational structure that is not captured by displayed peptides. Profiling conformational protein interactions at proteome scale has traditionally relied on protein microarray technologies. Protein microarrays, however, tend to suffer from high per-assay cost, and a myriad of technical artifacts, including those associated with the high throughput expression and purification of proteins, the spotting of proteins onto a solid support, the drying and rehydration of arrayed proteins, and the slide-scanning fluorescence imaging-based readout.(*5, 6*) Alternative approaches to protein microarray production and storage have been developed (e.g. Nucleic Acid-Programmable Protein Array, NAPPA(*7*) or single-molecule PCR-linked in vitro expression, SIMPLEX(*8*)), but a robust, scalable, and cost-effective technology has been lacking.

To overcome the limitations associated with array-based profiling of full-length proteins, we previously established a methodology called ParalleL Analysis of Translated Open reading frames (PLATO), which utilizes ribosome display of open reading frame (ORF) libraries.(*9*) Ribosome display relies on *in vitro* translation of mRNAs that lack stop codons, stalling ribosomes at the ends of mRNA molecules in a complex with the nascent proteins they encode. PLATO suffers from several key limitations that have hindered its adoption. An ideal alternative is the covalent conjugation of proteins to short, amplifiable DNA barcodes. Indeed, individually prepared DNA-barcoded antibodies and proteins have been employed successfully in a myriad of applications, as reviewed recently by Liszczak and Muir.(*10*) One particularly attractive protein-DNA conjugation method involves the HaloTag system, which adapts a bacterial enzyme that forms an irreversible covalent bond with halogen-terminated alkane moieties.(*11*) Individual DNA-barcoded HaloTag fusion proteins have been shown to greatly enhance sensitivity and dynamic range of autoantibody detection, compared with traditional ELISA.(*12*) Scaling individual protein barcoding to entire ORFeome libraries would be immensely valuable, but formidable due to high cost and low throughput. Therefore, a self-assembly approach could provide a much more efficient path to library production.

Here we describe a novel molecular display technology, Molecular Indexing of Proteins by Self Assembly (MIPSA), which overcomes key disadvantages of PLATO and other full-length protein array technologies. MIPSA produces libraries of soluble full-length proteins, each uniquely identifiable via covalent conjugation to a DNA barcode, flanked by universal PCR primer binding sequences. Barcodes are introduced near the 5’ end of transcribed mRNA sequences, upstream of the ribosome binding site (RBS). Reverse transcription (RT) of the 5’ end of *in vitro* transcribed RNA (IVT-RNA) creates a cDNA barcode, which is linked to a haloalkane-labeled RT primer. An N-terminal HaloTag fusion protein is encoded downstream of the RBS, such that *in vitro* translation results in the intra-complex (“*cis*”), covalent coupling of the cDNA barcode to the HaloTag and its downstream open reading frame (ORF) encoded protein product. The resulting library of uniquely indexed full-length proteins can be used for inexpensive proteome-wide interaction studies, such as unbiased autoantibody profiling. We demonstrate the utility of the platform by uncovering known and novel autoantibodies in the plasma of patients with severe COVID-19.

## Results

### Development of the MIPSA system

The MIPSA Gateway Destination vector contains the following key elements: a T7 RNA polymerase transcriptional start site, an isothermal unique clonal identifier (“UCI”) barcode sequence flanked by constant primer binding sites, a ribosome binding site (RBS), an N-terminal HaloTag fusion protein (891 nt), recombination sequences for ORF insertion, a stop codon, and a homing endonuclease site for plasmid linearization. A recombined ORF-containing pDEST-MIPSA plasmid is shown in **Fig. 1A**.

**Fig. 1.**
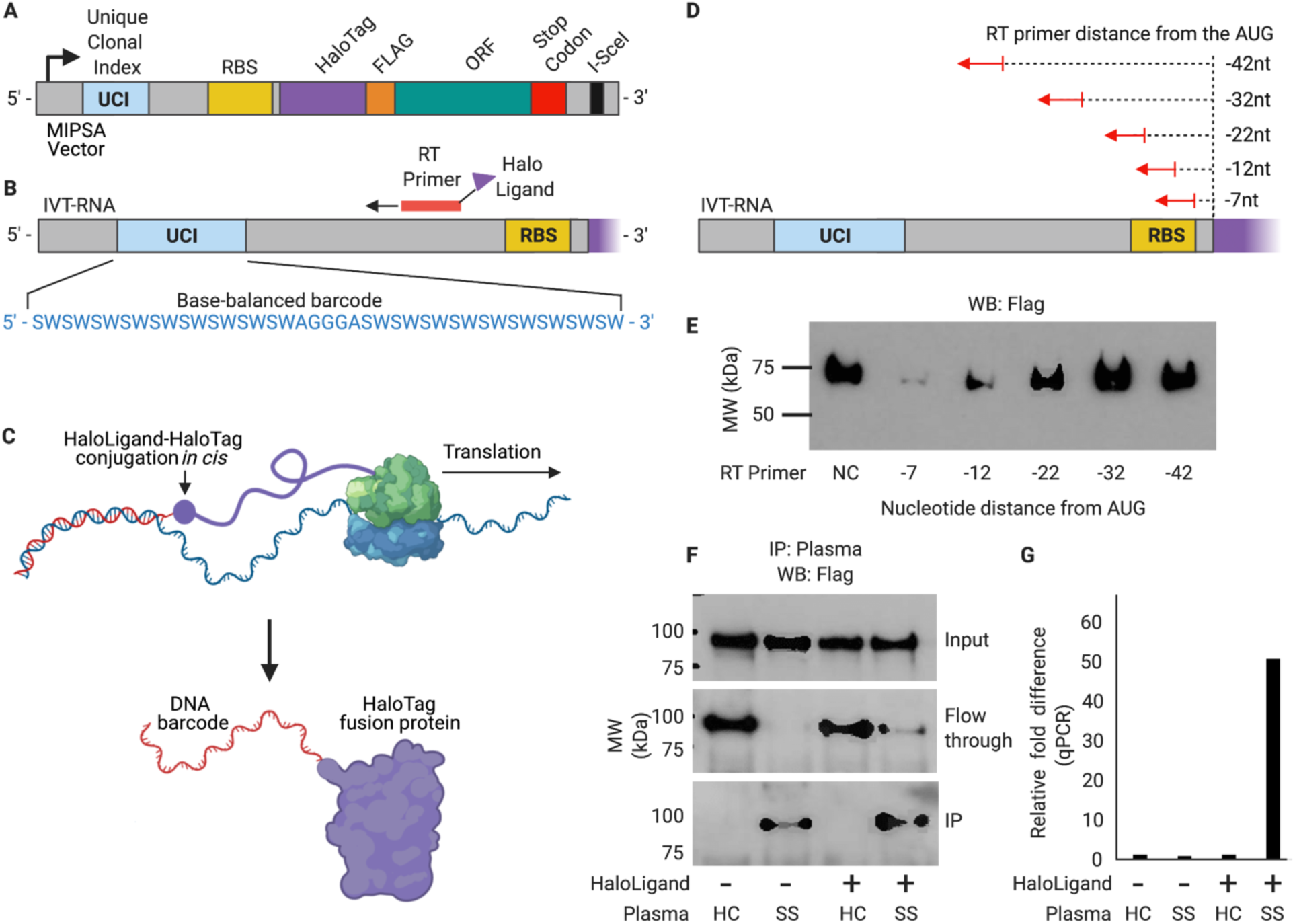
The MIPSA method. (**A**) Schematic of the recombined pDEST-MIPSA vector with key components highlighted: unique clonal index (UCI, blue), ribosome binding site (RBS, yellow), N-terminal HaloTag (purple), FLAG epitope (orange), open reading frame (ORF, green), stop codon (red) and the I-SceI restriction endonuclease site (black) for vector linearization. (**B**) Schematic showing *in vitro* transcribed (IVT) RNA from the vector template shown in **A**. Isothermal base-balanced UCI sequence: (SW)18-AGGGA-(SW)18. (**C**) Cell-free translation of the RNA-cDNA shown in **B**. HaloTag protein forms a covalent bond with the HaloLigand-conjugated UCI-containing cDNA *in cis* during translation. (**D**) RT primer positions tested for impact on translation. (**E**) α-FLAG western blot analysis of translation in presence of RT primers depicted in **D** (NC, negative control, no RT primer). (**F**) Western blot analysis of TRIM21 protein translated from RNA carrying the UCI-cDNA primed from the -32 position, either conjugated (+) or not (-) with the HaloLigand. Sjogren’s Syndrome, SS; Healthy Control, HC. (**G**) qPCR analysis of the IPed TRIM21 UCI. Fold-difference is by comparison with the HaloLigand (-) HC IP.

We first sought to establish a library of pDEST-MIPSA plasmids containing stochastic, isothermal UCIs located between the transcriptional start site and the ribosome binding site. A degenerate oligonucleotide pool was synthesized, comprising melting temperature (Tm) balanced sequences: (SW)18-AGGGA-(SW)18, where S represents an equal mix of C and G, while W represents an equal mix of A and T (**Fig. 1B**). We reasoned that this inexpensive pool of sequences would (i) provide sufficient complexity (2^36^ ∼ 7 x 10^10^) for unique ORF labeling, (ii) amplify without distortion, and (iii) serve as ORF-specific forward and reverse qPCR primer binding sites for measurement of individual UCIs of interest. The degenerate oligonucleotide pool was amplified by PCR, restriction cloned into the MIPSA destination vector, and transformed into *E. coli* (Methods). About 800,000 transformants were scraped off selection plates to obtain the pDEST-MIPSA UCI plasmid library. ORFs encoding the housekeeping gene glyceraldehyde-3-phosphate dehydrogenase (GAPDH) and a known autoantigen, tripartite motif containing-21 (TRIM21, commonly known as Ro52), were separately recombined into the pDEST-MIPSA UCI plasmid library and used in the following experiments. Individually barcoded GAPDH and TRIM21 clones were isolated and sequenced.

The MIPSA procedure involves RT of the stochastic barcode using a succinimidyl ester (O2)-haloalkane (HaloLigand)-conjugated RT primer. The bound RT primer should not interfere with the assembly of the *E. coli* ribosome and initiation of translation, but should be sufficiently proximal such that coupling of the HaloLigand-HaloTag-protein complex might hinder additional rounds of translation. We tested a series of RT primers that anneal at distances ranging from -42 nucleotides to -7 nucleotides (5’ to 3’) relative to the zero position of the AUG start codon (**Fig. 1D**). Based on the yield of protein product from mRNA saturated with primers at these varying locations, we selected the -32 position as it did not interfere with translation efficiency (**Fig. 1E**). In contrast, RT from primers located within 20 nucleotides of the RBS diminished or abolished protein translation. This result agrees with the estimated footprint of assembled 70S *E. coli* ribosomes, which have been shown to protect a minimum of 15 nucleotides of mRNA.(*13*)

We next assessed the ability of SuperScript IV to perform RT from a primer labeled with the HaloLigand at its 5’ end, and the ability of the HaloTag-TRIM21 protein to form a covalent bond with the HaloLigand-conjugated primer during the translation reaction. HaloLigand conjugation and purification followed Gu et al. (Methods, **Fig. S1**).(*14*) Either an unconjugated RT primer or a HaloLigand-conjugated RT primer was used for RT of the barcoded HaloTag-TRIM21 mRNA. The translation product was then immunoprecipitated (IPed) with plasma from a healthy donor or plasma from a TRIM21 (Ro52) autoantibody-positive patient with Sjogren’s Syndrome (SS). The SS plasma efficiently IPed the TRIM21 protein, regardless of RT primer conjugation, but only pulled down the TRIM21 cDNA UCI when the HaloLigand-conjugated primer was used in the RT reaction (**Fig. 1F-G**).

### Assessing cis versus trans UCI barcoding

While the previous experiment indicated that indeed the HaloLigand does not impede RT priming, and that the HaloTag can form a covalent bond with the HaloLigand during the translation reaction, it did not elucidate the amount of *cis* (intra-complex) versus *trans* (inter-complex) HaloTag-UCI conjugation (**Fig. S2**). Here, “intra-complex” is defined as conjugation to the UCI, which is bound to the same RNA encoding the protein. To measure the amount of *cis* and *trans* HaloTag-UCI conjugation, GAPDH and TRIM21 mRNAs were separately reverse transcribed (using HaloLigand-conjugated primer) and then either mixed 1:1 or kept separate for *in vitro* translation. As expected, translation of the mixture produced roughly equivalent amounts of each protein compared to the individual translations (**Fig. S3**). SS plasma specifically IPed TRIM21 protein regardless of translation condition (**Fig. S3**, IPed fraction). However, we noted that while the SS IPs contained high levels of the TRIM21 UCI, as intended, more of the GAPDH UCI was pulled down by the SS plasma compared to that by the HC plasma when the mRNA was mixed prior to translation. This indicates that indeed some *trans* barcoding occurs (**Fig. 2A**). We estimate that ∼50% of the protein is *cis*-barcoded, with the remaining 50% *trans*-barcoded protein equally represented by both UCIs. Thus, in this two-component system, 25% of the TRIM21 protein is conjugated to the GAPDH UCI.

**Fig. 2.**
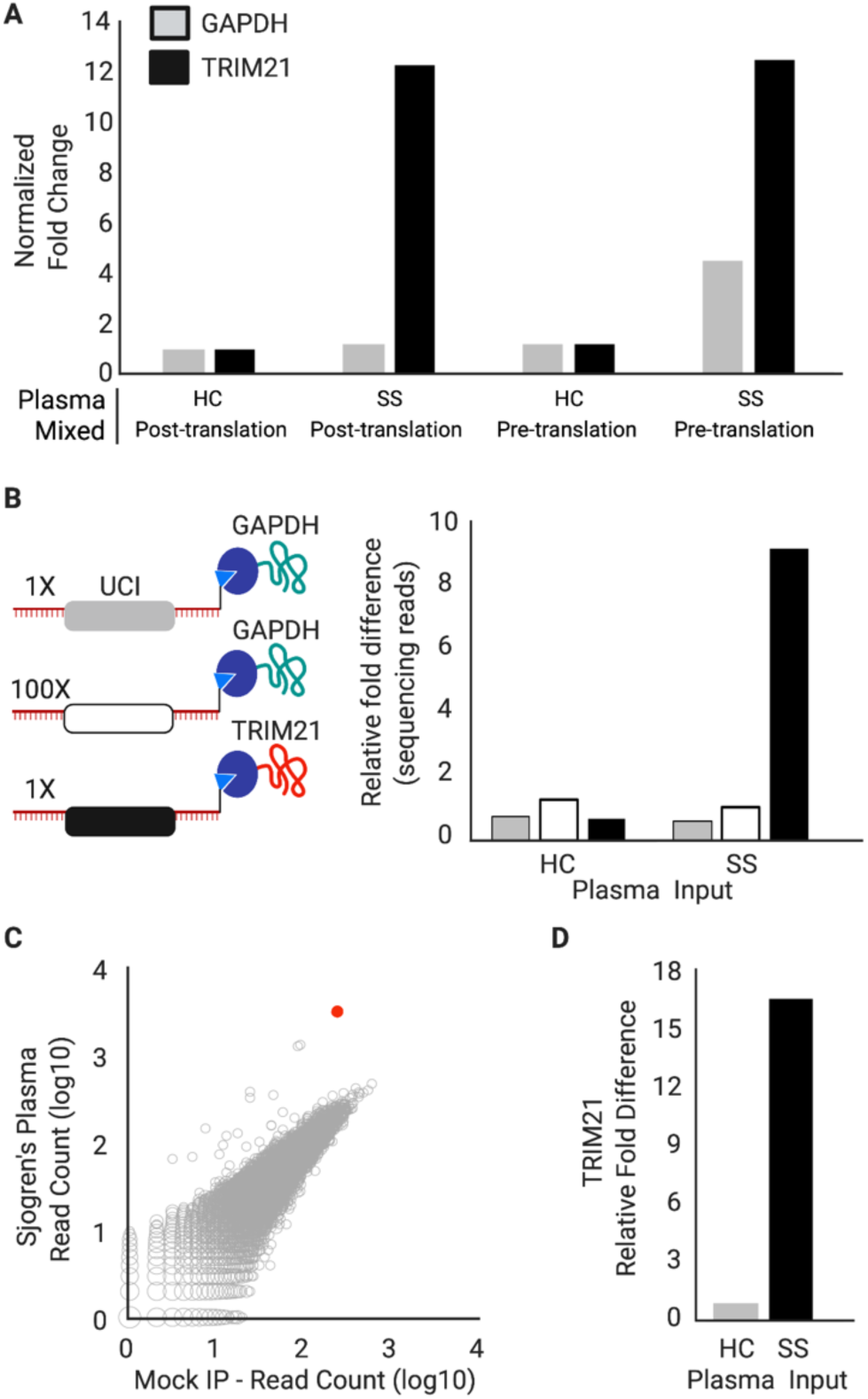
*Cis-* versus *trans*-UCI conjugation. (**A**) IVT-RNA encoding TRIM21 or GAPDH with their distinct UCI barcodes were translated before or after mixing at a 1:1 ratio. qPCR analysis of the IPs using UCI-specific primers, reported as fold-change versus IP with HC plasma, when the IVT-RNA was mixed post-translation. (**B**) IVT-RNA encoding TRIM21 (black UCI) and GAPDH (gray UCI) were mixed 1:1 into a background of 100-fold excess GAPDH (white UCI) and then translated. Sequencing analysis of the IPs, reported as fold-change versus the HC IP of the 100x GAPDH. (**C**) hORFeome MIPSA library containing spiked-in TRIM21, IPed with SS plasma and compared to average of 8 mock IPs (no plasma input). The TRIM21 UCI is shown in red. (**D**) Relative fold difference of TRIM21 UCI in SS vs HC IPs, determined by sequencing.

In the setting of a complex library, even if ∼50% of the protein is *trans* barcoded, this side product is uniformly distributed across all members of the library. We tested this hypothesis using a model MIPSA library composed of 100-fold excess of a second GAPDH clone, which was combined with a 1:1 mixture of the first GAPDH and TRIM21 clones (**Fig. 2B**). We additionally developed a sequencing workflow utilizing a PCR spike-in sequence for absolute quantification of each UCI. IP with SS plasma resulted in the specific IP of the TRIM21-UCI, with negligible *trans*-coupled GAPDH-UCI detected (**Fig. 2B**). Using the spiked-in sequence for absolute quantification, and assuming of 100% of the input TRIM21 protein in the IP fraction, we calculated a *cis* coupling efficiency of about 0.2% (i.e. 0.2% of input TRIM21 RNA molecules were converted into the intended *cis* UCI-coupled TRIM21 proteins).

### Establishing and deconvoluting a stochastically barcoded human ORFeome MIPSA library

The sequence-verified human ORFeome (hORFeome) v8.1 is composed of 12,680 clonal ORFs mapping to 11,437 genes in pDONR223.(*15*) Five subpools of the library were created, each composed of ∼2,500 similarly sized ORFs. Each of the five subpools was separately recombined into the pDEST-MIPSA UCI plasmid library and transformed to obtain ∼10-fold ORF coverage (∼25,000 clones per subpool). Each subpool was assessed via Bioanalyzer electrophoresis, sequencing of ∼20 colonies, and Illumina sequencing of the superpool. The TRIM21 plasmid was spiked into the superpooled hORFeome library at 1:10,000 – comparable to a typical library member. The SS IP experiment was then performed on the hORFeome MIPSA library, using sequencing as a readout. The reads from all barcodes in the library, including the spiked-in TRIM21, are shown in **Fig. 2C**. The SS autoantibody-dependent enrichment of TRIM21 (17-fold) was similar to the model system (**Fig. 2D**). Assuming the coupling efficiencies derived earlier, we estimate that about 6×10^5^ correctly *cis*-coupled TRIM21 molecules (and thus each library member on average) was input to the IP reaction.

Next, we established a system for creating a UCI-ORF lookup dictionary, using tagmentation and sequencing (**Fig. 3A**). Sequencing the 5’ 50 nt of the ORF inserts detected 11,076 of the 11,887 unique 5’ 50 nt sequences. Of the 153,161 unique barcodes detected, 82.9% (126, 975) were found to be associated with a single ORF (“monoclonal”). Each ORF was uniquely associated with a median of 9 (ranging from 0 to 123) UCIs (**Fig. 3B**). Aggregating the reads corresponding to each ORF, over 99% of the represented ORFs were present within a 10-fold difference of the median ORF abundance (**Fig. 3C**). Taken together, these data indicated that we established a uniform library of 11,076 stochastically indexed human ORFs, and sufficiently defined a lookup dictionary for downstream analyses. **Fig. 3D** shows a SS IP versus mock IP, but with the 47 dictionary-decoded GAPDH UCIs (corresponding to two GAPDH isoforms present in the hORFeome library) appearing along the y=x diagonal as expected.

**Fig. 3.**
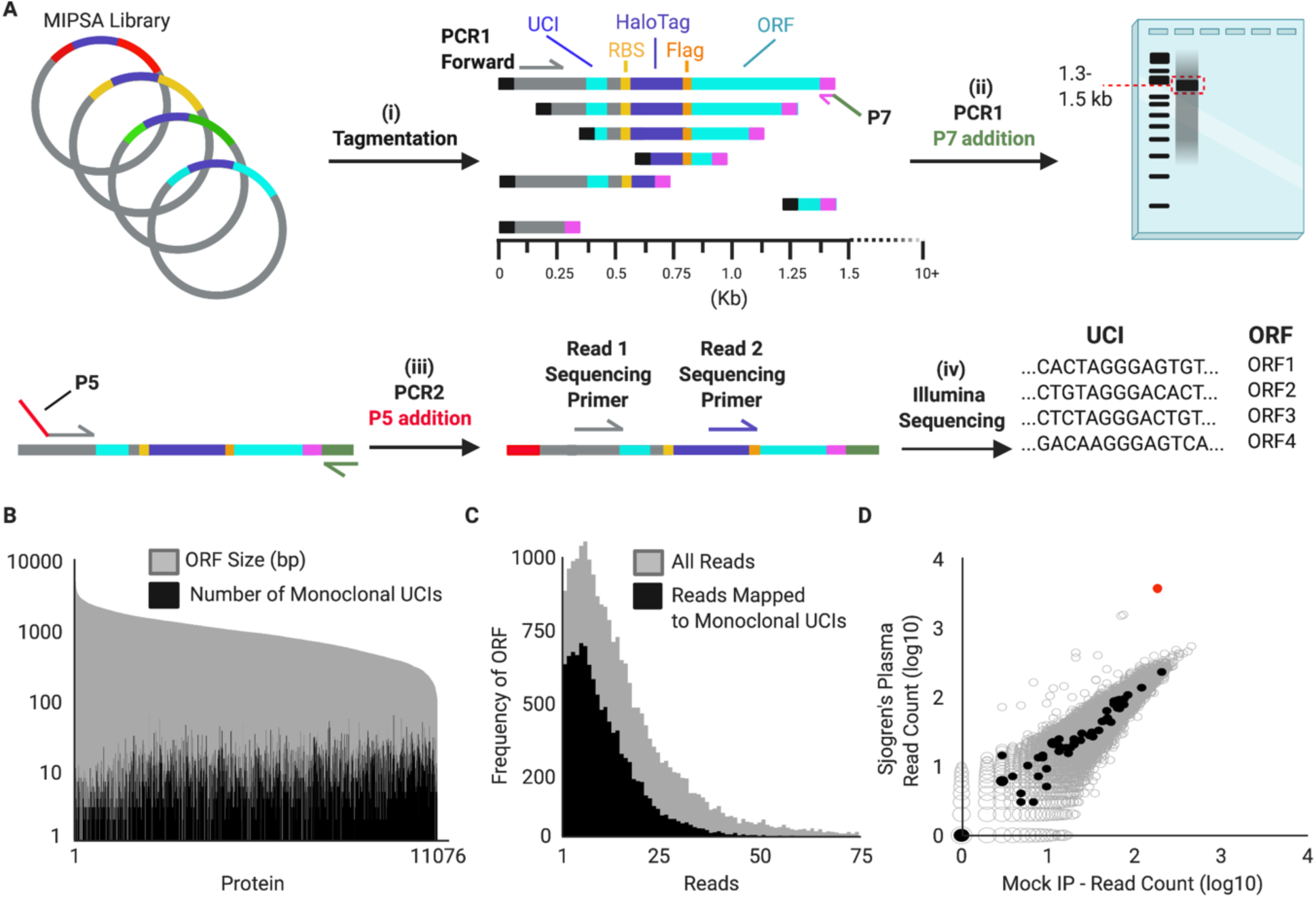
Defining the UCI-ORF dictionary. (**A**) Tagmentation randomly inserts adapters into the MIPSA vector library. Utilizing a PCR1 forward primer and the reverse primer of the tagmentation-inserted adapter, DNA fragments are amplified and size selected to be ∼1.5 kb, which captures the 5’ terminus of the ORF. These fragments are amplified with a P5-containing PCR2 forward primer and a P7 reverse primer. Illumina sequencing is used to read the UCI and the ORF from the same fragment, thus enabling their association in the dictionary. (**B**) The number of uniquely-associated UCIs is shown for each member of pDEST-MIPSA hORFeome library, superimposed on the length of the ORF. (**C**) Distribution of reads associated with each ORF, both total reads and UCI-matched reads. (**D**) IP of hORFeome MIPSA library using Sjogren’s Syndrome (SS) plasma is compared to the average of 8 mock IPs. Sequencing reads of each UCI are plotted. UCIs associated with the two library GAPDH isoforms (filled black) and spiked-in TRIM21 (red) are indicated.

### Unbiased MIPSA analysis of autoantibodies associated with severe COVID-19

Several recent reports have described elevated autoantibody reactivities in patients with severe COVID-19.(*16–20*) We therefore used MIPSA with the human ORFeome library for unbiased identification of autoreactivities in the plasma of 55 severe COVID-19 patients. For comparison, we used MIPSA to detect autoreactivities in plasma from 10 healthy donors and 10 COVID-19 convalescent plasma donors who had not been hospitalized (**Table S1**). Each sample was compared to a set of 8 “mock IPs”, which contained all reaction components except for plasma. Comparison to mock IPs accounts for bias in the library and background binding. Importantly, the informatic pipeline used to detect antibody-dependent reactivity (Methods) yielded a median of 5 (ranging from 2 to 9) false positive UCI hits per mock IP. IPs using plasma from severe COVID-19 patients, however, yielded a mean of 132 reactive UCIs, significantly more than the mean of 93 reactive UCIs among the controls (p = 0.018, t-test). Collapsing UCIs to their corresponding proteins yielded a mean of 83 reactive proteins among severe COVID-19 patients, which was significantly more than the mean of 63 reactive proteins among controls (**Fig. 4A**, p = 0.019, t-test).

**Fig. 4.**
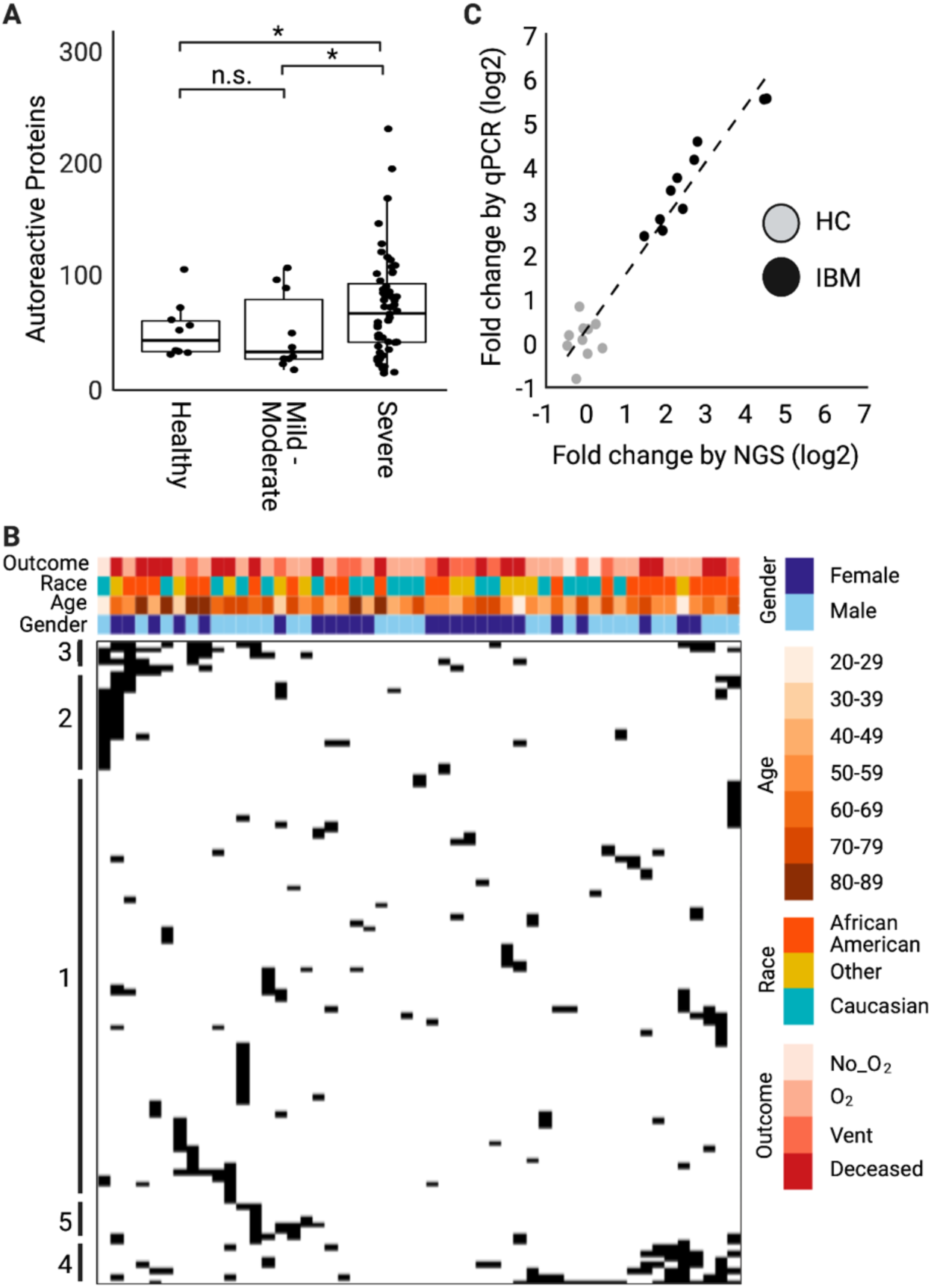
MIPSA analysis of autoantibodies in severe COVID-19. (**A**) Boxplots showing total numbers of reactive proteins in plasma from healthy controls, mild-moderate COVID-19 patients, or severe COVID-19 patients. * indicates p<0.05 using t-test to compare means. (**B**) Hierarchal clustering of all proteins represented by at least 2 reactive UCIs in at least 1 severe COVID-19 plasma, but not more than 1 control (healthy or mild-moderate COVID-19 plasma). (**C**) MIPSA analysis of autoantibodies in 10 Inclusion Body Myositis (IBM) patients and 10 healthy controls (HCs), using the hORFeome library. Fold change of IPed 5’-nucleotidase, cytosolic 1A (NT5C1A), measured by UCI-qPCR (relative to average of 10 HCs) or sequencing (relative to mock IPs).

We next examined proteins in the severe COVID-19 IPs that had at least two reactive UCIs (in the same IP), which were reactive in at least one severe patient, and that were not reactive in more than one control (healthy or mild/moderate convalescent plasma). Proteins were excluded if they were reactive in a single severe patient and a single control. The 103 proteins that met these criteria are shown in the clustered heatmap of **Fig. 4B**. Fifty one of the 55 severe COVID-19 patients exhibited reactivity to at least one of these proteins. We noted co-occurring protein reactivities in multiple individuals, the vast majority of which lack homology by protein sequence alignment. **Table S2** provides summary statistics about these reactive proteins, including whether they are previously defined autoantigens according to the human autoantigen database AAgAtlas 1.0.(*21*) **Data S1** provides the patient versus UCI-level data used to construct the heatmap.

One notable autoreactivity cluster (**Table S2,** cluster #5) includes 5’-nucleotidase, cytosolic 1A (NT5C1A), which is highly expressed in skeletal muscle and is the most well-characterized autoantibody target in inclusion body myositis (IBM). Multiple UCIs linked to NT5C1A were significantly increased in 3 of the 55 severe COVID-19 patients (5.5%). NT5C1A autoantibodies have been reported in up to 70% of IBM patients (*1*), in ∼20% of Sjogren’s Syndrome (SS) patients, and in up to ∼5% of healthy donors.(*22*) The prevalence of NT5C1A reactivity in the severe COVID-19 cohort is therefore not necessarily elevated. However, we wondered whether MIPSA would be able to reliably distinguish between healthy donor and IBM plasma based on NT5C1A reactivity. We tested plasma from 10 healthy donors and 10 IBM patients, the latter of whom were selected based on NT5C1A seropositivity determined by PhIP-Seq.(*1*) The clear separation of patients from controls in this independent cohort suggests that MIPSA may indeed have utility in clinical diagnostic testing using either UCI-specific qPCR or library sequencing, which were tightly correlated readouts (**Fig. 4C**).

### Type I and type III interferon-neutralizing autoantibodies in severe COVID-19 patients

Neutralizing autoantibodies targeting type I interferons alpha (IFN-α) and omega (IFN-ω) have been associated with severe COVID-19.(*17, 23, 24*) All type I interferons except IFN-α16 are represented in the human MIPSA library and dictionary. However, IFN-α4, IFN-α17, and IFN-α21 are indistinguishable by the first 50 nucleotides of their encoding ORF sequences. Two of the severe COVID-19 patients in this cohort (3.6%) exhibited dramatic IFN-α autoreactivity (43 and 41 IFN-α UCIs, across 10 distinct ORFs, along with 5 and 2 IFN-ω UCIs, **Fig. 5A-B**). The extensive co-reactivity of these proteins is likely attributable to their sequence homology (**Fig. S4**). By requiring at least 2 IFN UCIs to be considered positive, we identified two additional severe COVID-19 plasma (P3-P4) with lower levels of IFN-α reactivity, each with only 2 reactive IFN-α UCIs. Interestingly, one additional plasma (P5) precipitated five UCIs from the type III interferon IFN-λ3, but no UCI from any type I or II interferon (**Fig. 5C-D**). None of the healthy or non-hospitalized COVID-19 controls were positive for 2 or more interferon UCIs.

**Fig. 5.**
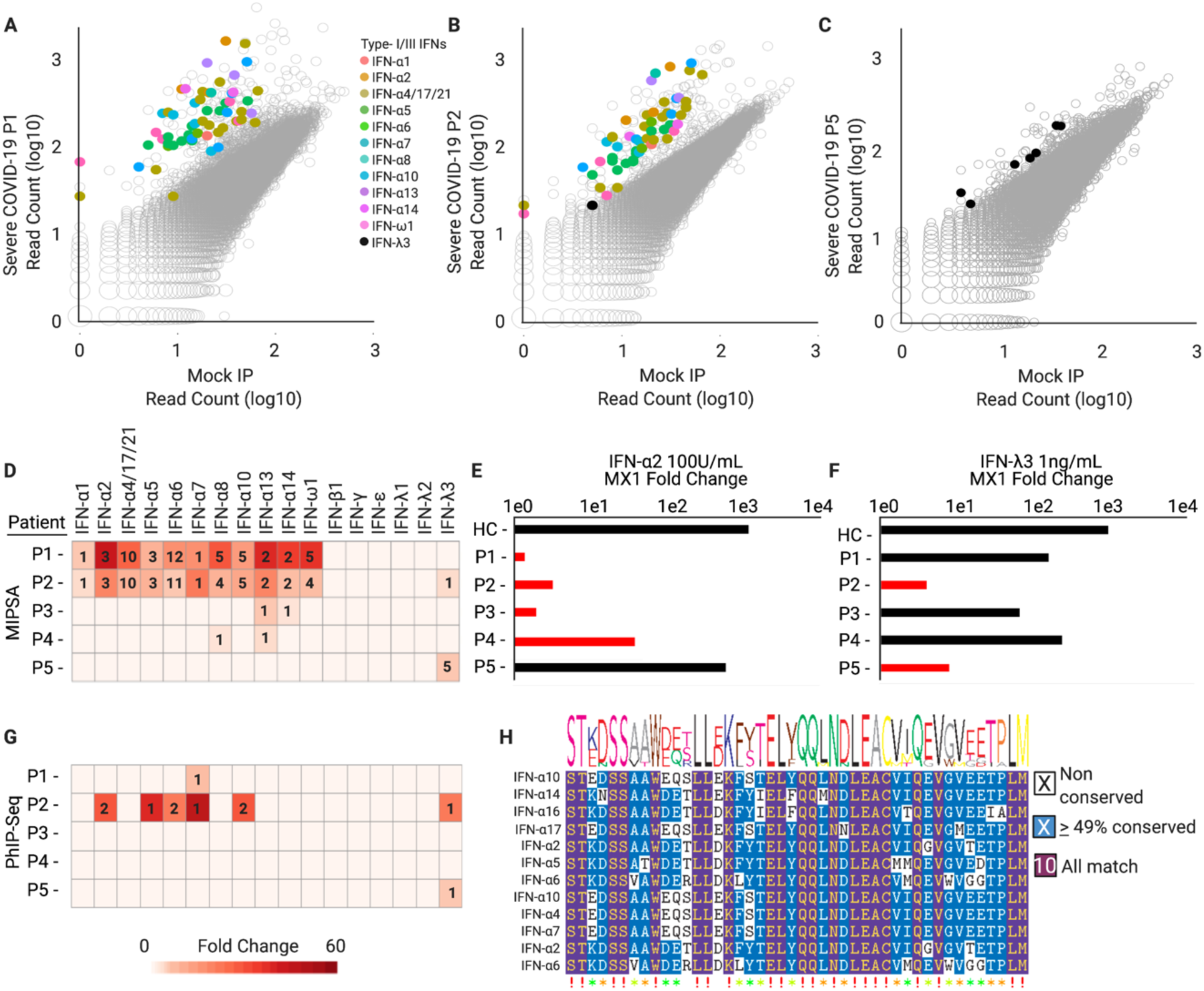
MIPSA detects known and novel neutralizing interferon autoantibodies. (**A-C**) Scatterplots highlighting reactive interferon UCIs for three severe COVID-19 patients. (**D**) Summary of interferon reactivity detected in 5 of 55 individuals with severe COVID-19. Hits fold-change values (cell color) and the number of reactive UCIs (number in cell) are provided. (**E, F**) Recombinant interferon alpha 2 (IFN-α2) or interferon lambda 3 (IFN-λ3) neutralizing activity of the same patients shown in **D**. Plasma were pre-incubated with 100 U/ml of IFN-α2 or 1 ng/ml of IFN-λ3 prior to incubation with A549 cells. Fold changes of the interferon stimulated gene, MX1, were calculated by RT-qPCR relative to unstimulated cells. GAPDH was used as a housekeeping control gene for normalization. Red bars indicate which samples are predicted by MIPSA to have neutralizing activity for each interferon. (**G**) PhIP-Seq analysis of interferon autoantibodies in the 5 patients of **D** (row and column orders maintained). Hits fold-change values (cell color) and the number of reactive peptides (number in cell) are provided. (**H**) Epitopefindr analysis of the PhIP-Seq reactive type I interferon 90-aa peptides.

Incubation of A549 human adenocarcinomatous lung epithelial cells with 100 U/ml IFN-α or 1 ng/ml of IFN-λ3 for 4 hours in serum-free medium resulted in a robust upregulation of the IFN-response gene MX1 by ∼1,000-fold and ∼100-fold, respectively. Pre-incubation of the IFN-α2 with plasma P1, P2, or P3 completely abolished the A549 interferon response (**Fig. 5E**). The plasma with the weakest IFN-α reactivity by MIPSA (P4) partially neutralized the cytokine. Neither HC nor P5 plasma had any effect on the response of A549 cells to IFN-α2. However, pre-incubation of the IFN-λ3 with the MIPSA-reactive plasma, P2 and P5, neutralized the cytokine (**Fig. 5F**). None of the other plasma (HC, P1, P3, or P4) had any effect on the response of A549 cells to IFN-λ3. In summary, antibody profiling of this severe COVID-19 cohort identified strongly neutralizing IFN-α autoantibodies in 5.5% of patients and strongly neutralizing IFN-λ3 autoantibodies in 3.6% of patients, with a single patient (1.8%) harboring both autoreactivities.

We then asked if PhIP-Seq with a 90-aa human peptidome library(*25*) might also detect interferon antibodies in this cohort. PhIP-Seq detected IFN-α reactivity in plasma from P1 and P2, although to a much lesser extent (**Fig. 5G**). The two weaker IFN-α reactivities detected by MIPSA in the plasma of P3 and P4 were both missed by PhIP-Seq. PhIP-Seq identified a single additional weakly IFN-α reactive sample, which was negative by MIPSA (not shown). Detection of type III interferon autoreactivity (directed exclusively at IFN-λ3) agreed perfectly between the two technologies. PhIP-Seq data was used to narrow the location of a dominant epitope in these type I and type III interferon autoantigens (**Fig. 5H** for IFN-α, amino acid position 45-135 for IFN-λ3).

We next wondered about the prevalence of the IFN-λ3 autoreactivity in the general population, and whether it might be increased among patients with severe COVID-19. PhIP-Seq was used to profile the plasma of 423 healthy controls, none of whom were found to have detectable IFN-λ3 autoreactivity. These data suggest that IFN-λ3 autoreactivity may be more frequent among individuals with severe COVID-19. This is the first report describing neutralizing IFN-λ3 autoantibodies, and therefore proposes a potentially novel pathogenic mechanism contributing to life-threatening COVID-19 in a subset of patients.

## Discussion

Here we present a novel molecular display technology for full length proteins, which provides key advantages over protein microarrays, PLATO, and alternative techniques. MIPSA utilizes self-assembly to produce a library of proteins, linked to relatively short (158 nt) single stranded DNA barcodes via the 25 kDa HaloTag domain. This compact barcoding approach will likely have numerous applications not accessible to alternative display formats with bulky linkage cargos (e.g. yeast, bacteria, viruses, phage, ribosomes, mRNAs). Indeed, individually conjugating minimal DNA barcodes to proteins, especially antibodies and antigens, has already proven useful in several contexts, including CITE-Seq,(*26*) LIBRA-seq,(*27*) and related methodologies.(*23, 28*) At proteome scale, MIPSA enables unbiased analyses of protein-antibody, protein-protein, and protein-small molecule interactions, as well as studies of post-translational modification, such as hapten modification studies or protease activity profiling, for example. Key advantages of MIPSA include its high throughput, low cost, simple sequencing library preparation, and stability of the protein-DNA complexes (important for both manipulation and storage of display libraries). Importantly, MIPSA can be immediately adopted by standard molecular biology laboratories, since it does not require specialized training or instrumentation, simply access to a high throughput DNA sequencing instrument or facility.

### Complementarity of MIPSA and PhIP-Seq

Display technologies frequently complement one another, but may not be amenable to routine use in concert. MIPSA is more likely than PhIP-Seq to detect antibodies directed at conformational epitopes on proteins expressed well *in vitro*. This was exemplified by the robust detection of interferon alpha autoantibodies via MIPSA, which were less sensitively detected via PhIP-Seq. PhIP-Seq, on the other hand, is more likely to detect antibodies directed at less conformational epitopes contained within proteins that are either absent from an ORFeome library or cannot be expressed well in cell free lysate. Because MIPSA and PhIP-Seq naturally complement one another in these ways, we designed the MIPSA UCI amplification primers to be the same as those we have used for PhIP-Seq. Since the UCI-protein complex is stable – even in bacterial phage lysate – MIPSA and PhIP-Seq can readily be performed together in a single reaction, using a single set of amplification and sequencing primers. The compatibility of these two display modalities will therefore lower the barrier to leveraging their synergy.

### Variations of the MIPSA system

A key aspect of MIPSA involves the conjugation of a protein to its associated UCI in *cis*, compared to another library member’s UCI in *trans*. Here we have utilized covalent conjugation via the HaloTag/HaloLigand system, but there are others that could work as well. For instance, the SNAP-tag (a 20 kDa mutant of the DNA repair protein O6-alkylguanine-DNA alkyltransferase) forms a covalent bond with benzylguanine (BG) derivatives.(*29*) BG could thus be used to label the RT primer in place of the HaloLigand. A mutant derivative of the SNAP-tag, the CLIP-tag, binds O2-benzylcytosine derivatives, which could also be adapted to MIPSA.(*30*)

The rate of fusion tag maturation and ligand binding is critical to the relative yield of *cis* versus *trans* UCI conjugation. A study by Samelson et al. determined that the rate of HaloTag protein production is about fourfold higher than the rate of HaloTag functional maturation.(*31*) Considering a typical protein size is <1,000 amino acids in the ORFeome library, these data predict that most proteins would be released from the ribosome before HaloTag maturation and thus before *cis* HaloLigand conjugation could occur, thereby favoring unwanted *trans* barcoding. However, we observed ∼50% of protein-UCI conjugates are formed *in cis*, thereby enabling sufficient assay performance in the setting of a complex library. During optimization experiments, we found the rate of *cis* barcoding to be slightly improved by excluding release factors from the translation mix, which stalls ribosomes on their stop codons and allows HaloTag maturation to continue in proximity to its UCI. Alternative approaches to promote controlled ribosomal stalling could include stop codon removal/suppression or use of a dominant negative release factor. Ribosome release could then be induced via addition of the chain terminator puromycin.

Since UCI cDNAs are formed on the 5’ UTR of the IVT-RNA, eukaryotic ribosomes would be unable to scan from the 5’ cap to the initiating Kozak sequence. The MIPSA system described here is therefore incompatible with cap-dependent cell-free translation systems. In case cap-dependent translation is required, however, two alternative methods could be developed. First, the current 5’ UCI system could be used if an internal ribosome entry site (IRES) were to be placed between the RT primer and the Kozak sequence. Second, the UCI could instead be situated at the 3’ end of the RNA, provided that the RT was prevented from extending into the ORF. Beyond cell-free translation, if either of these approaches were developed, RNA-cDNA hybrids could be transfected into living cells or tissues, where UCI-protein formation could take place *in situ*.

The ORF-associated UCIs can be embodied in a variety of ways. Here, we have stochastically assigned indexes to the human ORFeome at ∼10x representation. This approach has two main benefits, first being the low cost of the single degenerate oligonucleotide pool, and second being the multiple, independent pieces of evidence reported by the set of UCIs associated with each ORF. We have designed our library of stochastic barcodes to feature base-balanced sequences of uniform melting temperature, and thus more uniform PCR amplification efficiency. For simplicity, we have opted not to incorporate unique molecular identifiers (UMIs) into the primer, but this approach is compatible with MIPSA UCIs, and may potentially enhance quantitation. One disadvantage of stochastic indexing is the potential for ORF dropout, and thus the need for relatively high UCI representation; this increases the depth of sequencing required to quantify each UCI, and thus the overall per-sample cost. A second disadvantage is the requirement to construct a UCI-ORFeome matching dictionary. With short-read sequencing, we were unable to disambiguate a fraction of the library, comprised mostly of alternative isoforms. Using a long-read sequencing technology, such as PacBio or Oxford Nanopore Technologies, instead of or in addition to short-read sequencing technology could surmount incomplete disambiguation. As opposed to stochastic barcoding, individual UCI-ORF cloning is possible but costly and cumbersome. However, a smaller UCI set would provide the advantage of lower per-assay sequencing cost. We have previously developed a methodology to clone ORFeomes using Long Adapter Single Stranded Oligonucleotide (LASSO) probes.(*32*) Incorporating target-specific indexes into the capture probe library would result in uniquely indexed ORFs, without dramatically increasing the cost of the LASSO probe library. LASSO cloning of ORFeome libraries may therefore synergize with MIPSA-based applications.

### MIPSA readout via qPCR

A useful feature of appropriately designed UCIs is that they can also serve as qPCR readout probes. The degenerate UCIs that we have designed and used here (**Fig. 1B**) also comprise 18 nt base-balanced forward and reverse primer binding sites. The low cost and rapid turnaround time of a qPCR assay can thus be leveraged in combination with MIPSA. For example, incorporating assay quality control measures, such as the TRIM21 IP, can be used to qualify a set of samples prior to a more costly sequencing run. Troubleshooting and optimization can similarly be expedited by employing qPCR as a readout, rather than relying exclusively on NGS. qPCR testing of specific UCIs may theoretically also provide enhanced sensitivity compared to sequencing, and may be more amenable to analysis in a clinical setting.

### Autoantibodies detected in severe COVID-19 patients using MIPSA

The association between autoimmunity and severe COVID-19 disease is increasingly appreciated. In a cohort of 55 hospitalized individuals, we detected multiple established autoantibodies, including one that we have previously linked to inclusion body myositis.(*1*) We then tested the performance of MIPSA for detecting the NT5C1A autoantibody in a separate cohort of seropositive IBM patients and healthy controls. The results support future efforts in evaluating the clinical utility of MIPSA for standardized, comprehensive autoantibody testing. Such tests could utilize either single-plex qPCR or library sequencing as a readout.

While clusters of autoreactivities were observed in multiple individuals, it is not clear what role, if any, they may play in severe COVID-19. In larger scale studies, we expect that patterns of co-occurring reactivity, or reactivities towards proteins with related biological functions, may ultimately define new autoimmune syndromes associated with severe COVID-19. Neutralizing IFN-α/ω autoantibodies have been described in patients with severe COVID-19 disease and are presumed to be pathogenic.(*17*) These likely pre-existing autoantibodies, which occur very rarely in the general population, block restriction of viral replication in cell culture, and are thus likely to interfere with disease resolution. This discovery paved the way to identifying a subset of individuals at risk for life-threatening COVID-19, and proposed therapeutic use of interferon beta in this population of patients. In our study, MIPSA identified two individuals with extensive reactivity to the entire family of IFN-α cytokines. Indeed, plasma from both individuals, plus one individual with weaker IFN-α reactivity detected by MIPSA, robustly neutralized recombinant IFN-α2 in a lung adenocarcinomatous cell culture model. Unexpectedly, one individual in the cohort without IFN-α reactivity pulled down 5 IFN-λ3 UCIs. A second, IFN-α autoreactive individual, also pulled down a single IFN-λ3 UCI. The same autoreactivities were also detected using PhIP-Seq. Interestingly, neither MIPSA nor PhIP-Seq detected reactivity to IFN-λ2, despite their high degree of sequence homology (**Fig. S4**). We tested the IFN-λ3 neutralizing capacity of these patients’ plasma, observing near complete ablation of the cellular response to the recombinant cytokine (**Fig. 5F**). These data propose IFN-λ3 autoreactivity is a new, potentially pathogenic mechanism contributing to severe COVID-19 disease.

Type III IFNs (IFN-λ, also known as IL-28/29) are cytokines with potent anti-viral activities that act primarily at barrier sites. The IFN-λR1/IL-10RB heterodimeric receptor for IFN-λ is expressed on lung epithelial cells and is important for the innate response to viral infection. Mordstein et al., determined that in mice, IFN-λ diminished pathogenicity and suppressed replication of influenza viruses, respiratory syncytial virus, human metapneumovirus, and severe acute respiratory syndrome coronavirus (SARS-CoV-1).(*33*) It has been proposed that IFN-λ exerts much of its antiviral activity *in vivo* via stimulatory interactions with immune cells, rather than through induction of the antiviral cell state.(*34*) Importantly, IFN-λ has been found to robustly restrict SARS-CoV-2 replication in primary human bronchial epithelial cells(*35*), primary human airway epithelial cultures(*36*), and primary human intestinal epithelial cells(*37*). Collectively, these studies suggest multifaceted mechanisms by which neutralizing IFN-λ autoantibodies may exacerbate SARS-CoV-2 infections.

Casanova, et al. did not detect any type III IFN neutralizing antibodies among 101 individuals with type I IFN autoantibodies tested.(*17*) In our study, one of the three IFN-α autoreactive individuals (P2, a 22-year-old male) also harbored autoantibodies that neutralized IFN-λ3. It is possible that this co-reactivity is extremely rare and thus not represented in the Casanova cohort. Alternatively, it is possible that the differing assay conditions exhibit differing detection sensitivity. Whereas Casanova, et al. cultured A549 cells with IFN-λ3 at 50 ng/ml without plasma preincubation, we cultured A549 cells with IFN-λ3 at 1 ng/ml after pre-incubation with plasma for one hour. Their readout of STAT3 phosphorylation may also provide different detection sensitivity compared to the upregulation of MX1 expression. A larger study is needed to determine the true frequency of these reactivities in severe COVID-19 patients and matched controls. Here, we report strongly neutralizing IFN-α and IFN-λ3 autoantibodies in 3 (5.5%) and 2 (3.6%) individuals, respectively, of 55 patients with severe COVID-19. IFN-λ3 autoantibodies were not detected via PhIP-Seq in a larger cohort of 423 healthy controls collected prior to the pandemic.

Type III interferons have been proposed as a therapeutic modality for SARS-CoV-2 infection,(*36, 38–42*) and there are currently three ongoing clinical trials to test pegylated IFN-λ1 for efficacy in reducing morbidity and mortality associated with COVID-19 (ClinicalTrials.gov Identifiers: NCT04343976, NCT04534673, NCT04344600). One recently completed double-blind, placebo-controlled trial, NCT04354259, reported a significant reduction by 2.42 log copies per ml of SARS-CoV-2 at day 7 among mild to moderate COVID-19 patients in the outpatient setting (p=0·0041).(*43*) Future studies will determine whether anti-IFN-λ3 autoantibodies are pre-existing or arise in response to SARS-CoV-2 infection, and how often they also cross-neutralize IFN-λ1. Based on sequence alignment of IFN-λ1 and IFN-λ3 (∼29% homology, **Fig. S4**), however, cross-neutralization is expected to be rare, raising the possibility that patients with neutralizing IFN-λ3 autoantibodies may especially derive benefit from pegylated IFN-λ1 treatment.

## Conclusions

MIPSA is a new self-assembling protein display technology with key advantages over alternative approaches. It has properties that complement techniques like PhIP-Seq, and MIPSA ORFeome libraries can be conveniently screened in the same reactions with programmable phage display libraries. The MIPSA protocol presented here requires cap-independent, cell-free translation, but future adaptations may overcome this limitation. Applications for MIPSA-based studies include protein-protein, protein-antibody, and protein-small molecule interaction studies, as well as unbiased analyses of post-translational modifications. Here we used MIPSA to detect known autoantibodies and to discover neutralizing IFN-λ3 autoantibodies, among many other potentially pathogenic autoreactivities (**Table S2**), which may contribute to life-threatening COVID-19 pneumonia in a subset of at-risk individuals.

## Supporting information

DataS1

## Acknowledgements

We would like to thank Steve Elledge for generously providing the human ORFeome library and the 90-aa human peptidome T7 phage display library. We thank Rachel Green, Marco Catipovic, Allen Buskirk, Tim O’Donnell, Priya Duggal and Janet Markle for helpful discussions. We also thank Jodie Franklin and the Johns Hopkins Synthesis and Sequencing Core for HPLC purification of the HaloLigand conjugated RT-primer, as well as Linda Orzolek and Haiping Hao of the Johns Hopkins Transcriptomics and Deep Sequencing Core Facility.

The severe COVID-19 specimens utilized for this publication were part of the Johns Hopkins Biospecimen Repository, which is based on the contribution of many patients, research teams, and clinicians. Thanks to members of the NIH Vaccine Research Center for pre-pandemic sample collection: Barney Graham, Laura Novick, Joseph Casazza, Julie Ledgerwood, Uzma Sarwar, LeeJah Chang, Cynthia Starr Hendel, Lasonji Holman, Sarah Plummer, Pam Costner, Ingelise Gorden, Brenda Larkin, Floreliz Mendoza, Jamie Saudners, Kathy Zephir, Mary E Enama, Galina Yamshchikov, Iris Pittman, Pernell Williams.

All figures were created with BioRender.com

## Funding

Johns Hopkins University Provost Research Grant (HBL, ALC)

National Institute of General Medical Sciences grant R01GM127533 (HBL, BP)

National Heart, Lung, and Blood Institute of the National Institutes of Health grant K23HL151826 (EMB)

Sjogren’s Foundation and the Jerome L. Greene Foundation grants (ANB, HBL)

Vaccine Research Center, National Institute of Allergy and Infectious Diseases, National Institutes of Health intramural research program (MR)

## Author contributions

Conceptualization and experimental design: JJC, HBL Experiments: JJC, JG, PS, HJT, AW

Formal analysis: JJC, JG, PS, DRM, XAZ, WRM, YD, SJ, HBL

Funding acquisition: BP, HBL

Resources: ANB, MR, EMB, AART, IZ, JIS, AZR, ALC, TL, ALM

Visualization: JJC, JG, PS, DRM, XAZ, WRM

Writing: JJC, JG, PS, DRM, XAZ, HJT, AW, WRM, YD, SJ, LT, BP, ANB, MR, EMB, AART, IZ, JIS, AZR, ALC, TL, ALM, HBL

## Supplementary Materials

### Material and Methods

#### MIPSA Destination vector construction

The MIPSA vector was constructed using the Gateway pDEST15 vector as a backbone. A gBlock fragment (Integrated DNA Technologies) encoding the RBS, Kozak sequence, N-terminal HaloTag fusion protein, and FLAG tag, followed by an attR1 sequence was cloned into the parent plasmid. A stop codon and 150 bp poly(A) sequence was also added after attR2 site.

#### UCI barcode library construction

A 41 nt barcode oligo was generated within a gBlock Gene Fragment (Integrated DNA Technologies) with alternating mixed bases (S: G/C; W: A/T) to produce the following sequence: (SW)18-AGGGA-(SW)18. The sequences flanking the degenerate barcode incorporated the standard PhIP-Seq PCR1 and PCR2 primer binding sites.(*44*) 18 ng of the starting UCI library was used to run 40 cycles of PCR to amplify the library and incorporate BglII and PspxI restriction sites. The MIPSA vector and amplified UCI library were then digested with the restriction enzymes overnight, column purified, and ligated at 1:5 vector-to-insert ratio. The ligated MIPSA vector was used to transform electrocompetent One Shot ccdB 2 T1^R^ cells (Thermo Fisher Scientific). 6 transformation reactions yielded ∼800,000 colonies to produce the pDEST-MIPSA UCI library.

#### Human ORFeome recombination into the pDEST-MIPSA UCI plasmid library

150 ng of each pENTR-hORFeome subpool (L1-L5) was individually combined with 150 ng of the pDEST-MIPSA UCI library plasmid and 2 µl of Gateway LR Clonase II mix (Life Technologies) for a total reaction volume of 10 µl. The reaction was incubated overnight at 25°C. The entire reaction was transformed into 50 µl of One Shot OmniMAX 2 T1^R^ chemical competent *E. coli* (Life Technologies). In aggregate, the transformations yielded ∼120,000 colonies, which is ∼10-fold the complexity of the hORFeome v8.1. Colonies were collected and pooled by scraping, followed by purification of the barcoded pDEST-MIPSA-hORFeome plasmid DNA (human ORFeome MIPSA library) using the Qiagen Plasmid Midi Kit (Qiagen).

#### HaloLigand conjugation to RT oligo and HPLC purification

100 µg of a 5’ amine modified oligo HL-32_ad (**Table S3**) was incubated with 75 µl (17.85 µg/µl) of the HaloTag Succinimidyl Ester (O2) (Promega Corporation), the HaloLigand, in 0.1 M sodium borate buffer for 6 hours at room temperature following Gu, et al.(*14*) 3 M NaCl and ice-cold ethanol was added at 10% (v/v) and 250% (v/v), respectively, to the labeling reaction and incubated overnight at -80°C. The reaction was centrifuged for 30 minutes at 12,000 x g. The pellet was rinsed once in ice-cold 70% ethanol and air-dried for 10 minutes.

HaloLigand-conjugated RT primer was HPLC purified using a Brownlee Aquapore RP-300 7u, 100×4.6 mm column (Perkin Elmer) using a two-buffer gradient of 0–70% CH3CN/MeCN (100 mM triethylamine acetate to acetonitrile) over 70 minutes. Fractions corresponding to labeled oligo were collected and lyophilized (**Fig. S1**). Oligos were resuspended at 1 µM (15.4 ng/µl) and stored at -80°C.

#### MIPSA library IVT-RNA preparation

The human ORFeome MIPSA library plasmid (4 µg) was linearized with the I-SceI restriction endonuclease (New England Biolabs) overnight. The product was column-purified with the NucleoSpin Gel and PCR Clean Up kit (Macherey-Nagel). A 40 µl *in vitro* transcription reaction using the HiScribe T7 High Yield RNA Synthesis Kit (New England Biolabs) was utilized to transcribe 1 µg of the purified, linearized pDEST-MIPSA plasmid library. The product was diluted with 60 µl molecular biology grade water, and 1 µl of DNAse I was added. The reaction was incubated for another 15 minutes at 37°C. Then 50 µl of 1 M LiCl was added to the solution and incubated at -80°C overnight. A centrifuge was cooled to 4°C, and the RNA was spun at maximum speed for 30 minutes. The supernatant was removed, and the RNA pellet washed with 70% ethanol. The sample was spun down at 4°C for another 10 minutes, and the 70% ethanol removed. The pellet was dried at room temperature for 15 minutes, and subsequently resuspended in 100 µl water. To preserve the sample, 1 µl of 40 U/µl RNAseOUT Recombinant Ribonuclease Inhibitor (Life Technologies) was added.

#### MIPSA library IVT-RNA reverse transcription and translation

A reverse transcription reaction was prepared using SuperScript IV First-Strand Synthesis System (Life Technologies). First, 1 µl of 10 mM dNTPs, 1 µl of RNAseOUT (40 U/µl), 4.17 µl of the RNA library (1.5 µM), and 7.83 µl of the HaloLigand-conjugated RT primer (1 µM, **Table S3**) was combined for a single 14 µl reaction and incubated at 65°C for 5 minutes followed by a 2-minute incubation on ice. 4 µl of 5X RT buffer, 1 µl of 0.1 M DTT, and 1 µl of SuperScript IV RT Enzyme (200 U/µl) was added to the 14 µl reaction on ice and incubated for 20 minutes at 42°C. A single 20 µl RT reaction received 36 µl of RNAClean XP beads (Beckman Coulter) and was incubated at room temperature for 10 minutes. The beads were collected by magnet and washed five times with 70% ethanol. The beads were air-dried for 10 minutes at room temperature and resuspended in 7 µl of 5 mM Tris-HCl, pH 8.5. The product was analyzed with spectrophotometry to measure the RNA yield. A translation reaction was set up on ice using the PURExpress ΔRibosome Kit (New England Biolabs).(*45*) The reaction was modified such that the final concentration of ribosomes was 0.3 µM. For each 10 µl translation reaction, 4.57 µl of the RT reaction was added to 4 µl Solution A, 1.2 µl Factor Mix, and 0.23 µl ribosomes (13.3 µM). This reaction was incubated at 37°C for two hours, diluted to a total volume of 45 µl with 35 µl 1X PBS, and used immediately or stored at -80°C after addition of glycerol to a final concentration of 25% (v/v).

#### Immunoprecipitation of the translated MIPSA hORFeome library

5 µl of plasma, diluted 1:100 in PBS, is mixed with the 45 µl of diluted MIPSA library translation reaction (see above) and incubated overnight at 4°C with gentle agitation. For each IP, a mixture of 5 µl of Protein A Dynabeads and 5 µl of Protein G Dynabeads (Life Technologies) was washed 3 times in 2X their original volume with 1X PBS. The beads were then resuspended in 1X PBS at their original volume, and added to each IP. The antibody capture proceeded for 4 hours at 4°C. Beads were collected on a magnet and washed 3 times in 1X PBS, changing tubes or plates between washes. The beads were then collected and resuspended in a 20 µl PCR master mix containing the T7-Pep2_PCR1_F forward and the T7-Pep2_PCR1_R+ad_min reverse primers (**Table S3**) and Herculase-II (Agilent). PCR cycling was as follows: an initial denaturing and enzyme activation step at 95°C for 2 min, followed by 45 cycles of: 95°C for 20 s, 58°C for 30 s, and 72°C for 30 s. The final extension step was performed at 72°C for 3 minutes. Two microliters of the amplification product were used as input to a 20 µl dual-indexing PCR reaction for 10 cycles with the PhIP_PCR2_F forward and the Ad_min_BCX_P7 reverse primers. PCR cycling was as follows: an initial denaturing step at 95°C for 2 min, followed by 10 cycles of: 95°C for 20 s, 58°C for 30 s, and 72°C for 30 s. The final extension step was performed at 72°C for 3 min. i5/i7 indexed libraries were pooled and column purified (NucleoSpin columns, Takara). Libraries were sequenced on an Illumina NextSeq 500 using a 1×50 nt SE or 1×75 nt SE protocol. MIPSA_i5_NextSeq_SP and Standard_i7_SP primers were used for i5/i7 sequencing (**Table S3**) The output was demultiplexed using i5 and i7 without allowing any mismatches.

For quantification of MIPSA experiments by qPCR, the PCR1 product (above) was analyzed as follows. A 4.6 µl of 1:1,000 dilution of the PCR1 reaction was added to 5 µl of Brilliant III Ultra Fast 2X SYBR Green Mix (Agilent), 0.2 µl of 2 µM reference dye and 0.2 µl of 10 µM forward and reverse primer mix (specific to the target UCI). PCR cycling was as follows: an initial denaturing step at 95°C for 2 min, followed by 45 cycles of: 95°C for 20 s, 60°C for 30. Following completion of thermocycling, amplified products were subjected to melt-curve analysis. The qPCR primers for MIPSA immunoprecipitation experiments were: BT2_F and BT2_R for TRIM21, BG4_F and BG4_R for GAPDH, and NT5C1A_F and NT5C1A_R for NT5C1A (**Table S3**).

#### Plasma Samples

All samples were collected from subjects that met protocol eligibility criteria, as described below. All of the studies protected the rights and privacy of the study participants and were approved by their respective Intuitional Review Boards for original sample collection and subsequent analyses.

##### Pre-pandemic and healthy control plasma samples

All human samples were collected prior to 2017 at the National Institutes of Health (NIH) Clinical Center under the Vaccine Research Center’s (VRC)/National Institutes of Allergy and Infectious Diseases (NIAID)/NIH protocol “VRC 000: Screening Subjects for HIV Vaccine Research Studies” (NCT00031304) in compliance with NIAID IRB approved procedures.

##### COVID-19 Convalescent Plasma (CCP) from non-hospitalized patients

Eligible non-hospitalized CCP donors were contacted by study personnel, as previously described.(*46*) All donors were at least 18 years old and had a confirmed diagnosis of SARS-CoV-2 by detection of RNA in a nasopharyngeal swab sample. Basic demographic information (age, sex, hospitalization with COVID-19) was obtained from each donor; initial diagnosis of SARS-CoV-2 and the date of diagnosis were confirmed by medical chart review.

##### Severe COVID-19 plasma samples

The study cohort was defined as inpatients who had: 1) a confirmed RNA diagnosis of COVID-19 from a nasopharyngeal swab sample; 2) survival to death or discharge; and 3) remnant specimens in the Johns Hopkins COVID-19 Remnant Specimen Biorepository, an opportunity sample that includes 59% of Johns Hopkins Hospital COVID-19 patients and 66% of patients with length of stay ≥3 days.(*47, 48*) Patient outcomes were defined by the World Health Organization (WHO) COVID-19 disease severity scale. Samples from severe COVID-19 patients that were included in this study were obtained from 17 patients who died, 13 who recovered after being ventilated, 22 who required oxygen to recover, and 3 who recovered without supplementary oxygen. This study was approved by the JHU Institutional Review Board (IRB00248332, IRB00273516), with a waiver of consent because all specimens and clinical data were de-identified by the Core for Clinical Research Data Acquisition of the Johns Hopkins Institute for Clinical and Translational Research; the study team had no access to identifiable patient data.

##### Sjogren’s Syndrome and Inclusion body myositis (IBM) plasma samples

Sjogren’s syndrome samples were collected under protocol NA_00013201. All patients were >18 years old and gave informed consent. IBM patient samples were collected under protocol IRB00235256. All patients met ENMC 2011 diagnostic criteria(*49*) and provided informed consent.

#### Immunoblot analysis

Laemmli buffer containing 5% β-ME was added to samples, boiled for 5 min, and analyzed on NuPage 4-12% Bis-Tris polyacrylamide gels (Life Technologies). Following transfer to PVDF membranes, blots were blocked in 20 mM Tris-buffered saline, pH 7.6, containing 0.1% Tween 20 (TBST) and 5% (wt/vol) non-fat dry milk for 30 minutes at room temperature. Blots were subsequently incubated overnight at 4°C with primary anti-FLAG antibody (#F3165, MilliporeSigma) at 1:2,000 (v/v), followed by a 4-hour incubation at room temperature in anti-mouse IgG, HRP-linked secondary antibody (#7076, Cell Signaling) at 1:4,000 (v/v).

#### Construction of the UCI-ORF dictionary

The Nextera XT DNA Library Preparation kit (Illumina) was used for tagmentation of 150 ng of the pDEST-MIPSA hORFeome plasmid library to yield the optimal size distribution centered around 1.5 kb. Tagmented libraries were amplified using Herculase-II (Agilent) with T7-Pep2_PCR1_F forward and Nextera Index 1 Read primer. PCR cycling was as follows: an initial denaturing step at 95°C for 2 minutes, followed by 30 cycles of: 95°C for 20 s, 53.5°C for 30 s, 72°C for 30 s. A final extension step was performed at 72°C for 3 minutes. PCR reactions were run on a 1% agarose gel followed by excision of ∼1.5kb products and purification using the NucleoSpin Gel and PCR Clean-up columns (Macherey-Nagel). The purified product was then amplified for another 10 cycles with PhIP_PCR2_F forward and P7.2 reverse primers (see **Table S3** for list of primer sequences). The product was gel-purified and sequenced on a MiSeq (Illumina) using the T7-Pep2.2_SP_subA primer for read 1 and the MISEQ_MIPSA_R2 primer for read 2. Read 1 was 60 bp long to capture the UCIs. The first index read, I1, was substituted with a 50 bp read into the ORF. I2 was used to identify the i5 index for sample demultiplexing.

The hORFeome v8.1 DNA sequences were truncated to the first 50 nt, and the ORF names corresponding to non-unique sequences were concatenated with a “|” delimiter. The demultiplexed output of the 50 nt R2 (ORF) read from an Illumina MiSeq was aligned to the truncated human ORFeome v8.1 library using the Rbowtie2 package with the following parameters: options = “-a - -very-sensitive-local”.(*50*) The unique FASTQ identifiers were then used to extract corresponding sequences from the 60 bp R1 (UCI) read. Those sequences were then truncated using the 3’ anchor ACGATA, and sequences that did not have the anchor were removed. Additionally, any truncated R1 sequences that had fewer than 18 nucleotides were removed. The ORF sequences that still had a corresponding UCI post-filtering were retained using the FASTQ identifier. The names of ORFs that had the same UCI were concatenated with a “&” delimiter, and this final dictionary was used to generate a FASTA alignment file composed of ORF names and UCI sequences.

#### Informatic analysis of MIPSA sequencing data

Illumina output FASTQ files were truncated using the constant ACGAT anchor sequence following all UCI sequences. Next, perfect match alignment was used to map the truncated sequences to their linked ORFs via the UCI-ORF lookup dictionary. A counts matrix is constructed, in which rows correspond to individual UCIs and columns correspond to samples. We next used the edgeR software package(*51*) which, using a negative binomial model, compares the signal detected in each sample against a set of negative control (“mock”) IPs that were performed without plasma, to return a maximum likelihood fold-change estimate and a test statistic for each UCI in every sample, thus creating fold-change and -log10(p-value) matrices. Significantly enriched UCIs (“hits”) required a read count of at least 15, a p-value less than 0.001, and a fold change of at least 3. Hits fold-change matrices report the fold-change value for “hits” and report a “1” for UCIs that are not hits.

#### Protein sequence similarity

To evaluate sequence homology among proteins in the hORFeome v8.1 library, a blastp alignment was used to compare each protein sequence against all other library members (parameters: “-outfmt 6 -evalue 100 -max_hsps 1 -soft_masking false -word_size 7 - max_target_seqs 100000”). To evaluate sequence homology among reactive peptides in the human 90-aa phage display library, the epitopefindr(*52*) software was employed.

#### Phage ImmunoPrecipitation Sequencing (PhIP-Seq) analyses

PhIP-Seq was performed according to a previously published protocol.(*44*) Briefly, 0.2 µl of each plasma was individually mixed with the 90-aa human phage library and immunoprecipitated using protein A and protein G coated magnetic beads. A set of 6-8 mock immunoprecipitations (no plasma input) were run on each 96 well plate. Magnetic beads were resuspended in PCR master mix and subjected to thermocycling. A second PCR reaction was employed for sample barcoding. Amplicons were pooled and sequenced on an Illumina NextSeq 500 instrument using a 1×50 nt SE or 1×75 nt SE protocol. PhIP-Seq with the human library was used to characterize autoantibodies in a collection of plasma from healthy controls. For fair comparison to the severe COVID-19 cohort, we first determined the minimum sequencing depth that would have been required to detect the IFN-λ3 reactivity in both of the positive individuals. We then only considered the 423 data sets from the healthy cohort with sequencing depth greater than this minimum threshold. None of these 423 individuals were found to be reactive to any peptide from IFN-λ3.

#### Type I/III interferon neutralization assay

IFN-α2 (catalog no. 11100-1) and IFN-λ3 (catalog no. 5259-IL-025) were purchased from R&D Systems. 20 µl of plasma was incubated for 1 hour at room temperature with either 100 U/ml IFN-α2 or 1 ng/ml IFN-λ3, and 180 µl DMEM in a total volume of 200 µl before addition into 7.5×10^4^ A549 cells in 48-well tissue culture plates. After 4-hour incubation, the cells were washed with 1x PBS and cellular mRNA was extracted and purified using RNeasy Plus Mini Kit (Qiagen). 600 ng of extracted mRNA was reverse transcribed using the SuperScript III First-Strand Synthesis System (Life Technologies) and diluted 10-fold for qPCR analysis on a QuantStudio 6 Flex System (Applied Biosystems). PCR consisted of 95°C for 3 minutes, followed by 45 cycles of the following: 95°C for 15 seconds and 60°C for 30 seconds. MX1 expression was chosen as a measure of cell stimulation by the interferons, and the relative mRNA expression was normalized to GAPDH expression. The qPCR primers for GAPDH and MX1 were obtained from Integrated DNA Technologies (**Table S3**).

---

**Fig. S1.**
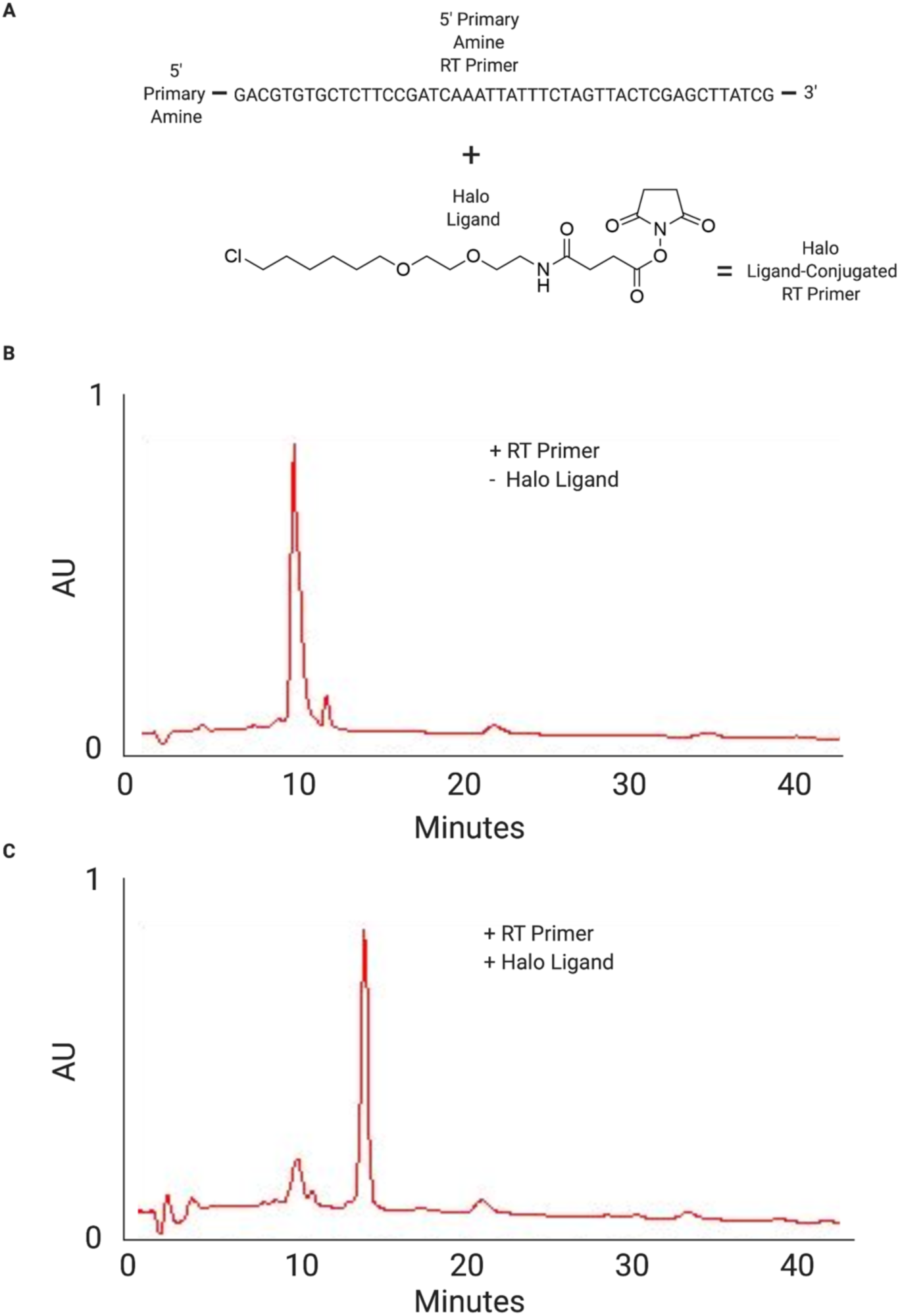
HaloLigand conjugation to the reverse transcription primer. (**A**) On the top is the oligonucleotide reverse transcription (RT) primer sequence modified with a 5’ primary amine. Below is the HaloLigand with a reactive succinimidyl ester group, separated by one ethylene glycol moiety (O2). The succinimidyl ester reacts with the primary amine to form an amide bond between the RT primer and the HaloLigand, resulting in the HaloLigand-conjugated RT primer. (**B**) HPLC chromatogram of the RT primer without the HaloLigand modification. (**C**) HPLC chromatogram of the RT primer with the HaloLigand modification after purification. The conjugated product is eluted off the column later due to its decreased hydrophobicity conferred by the modification.

**Fig. S2.**
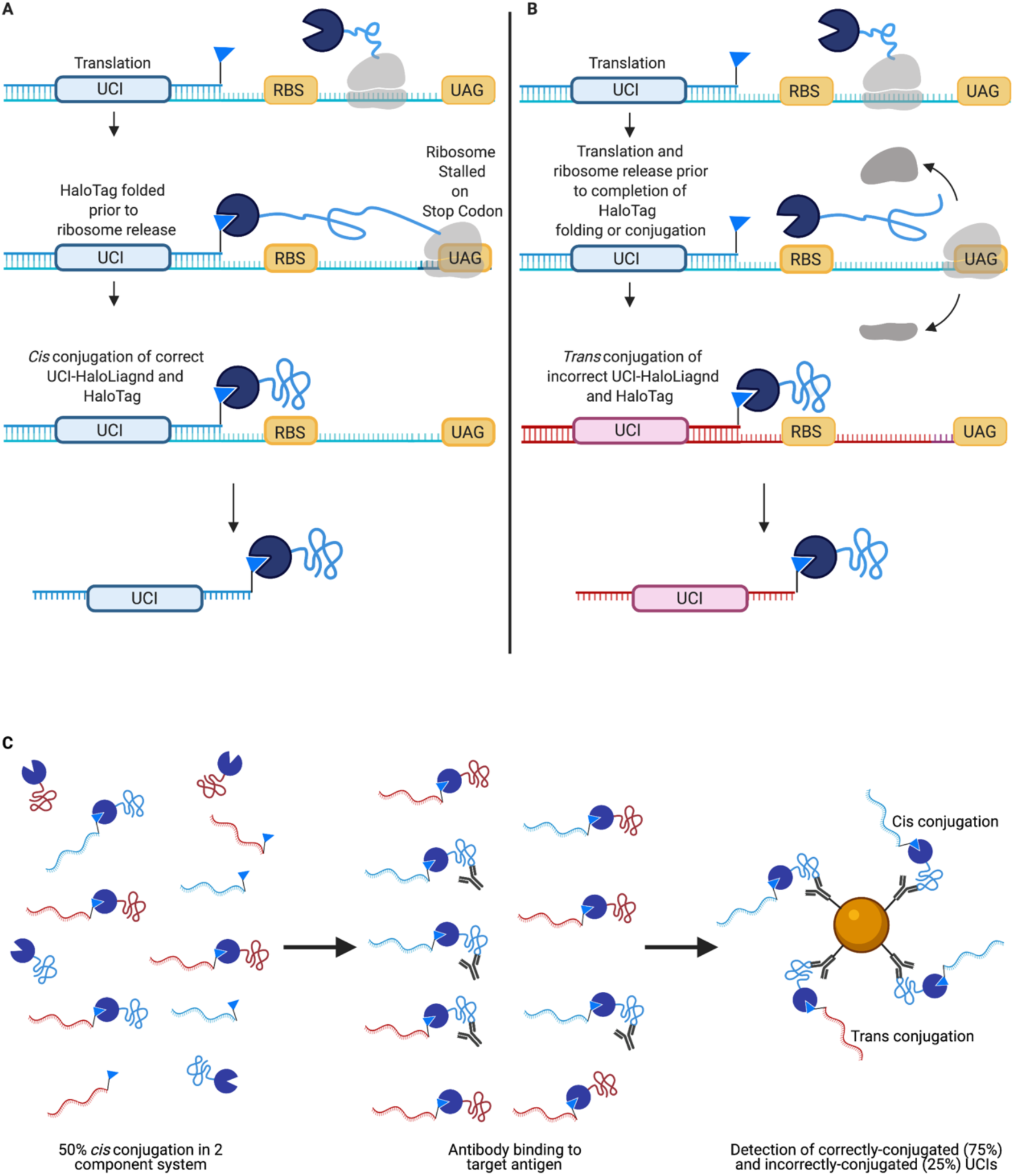
*Cis* versus *trans* UCI-ORF associations. Schematic of *cis* (**A**) versus *trans* (**B**) UCI-ORF conjugation during translation of a MIPSA IVT-RNA library. (**C**) Left panel: 50% *cis* conjugation composed of the correct protein-UCI associations (e.g. blue UCI with blue protein). Unconjugated proteins randomly associate with unconjugated UCIs (in *trans*). Middle panel: antibodies bind their target antigen. Right panel: the ratio of correctly to incorrectly IPed UCIs in this two-species experiment is 3:1 (75%:25%), similar to what was observed experimentally (Fig. 2A).

**Fig. S3.**
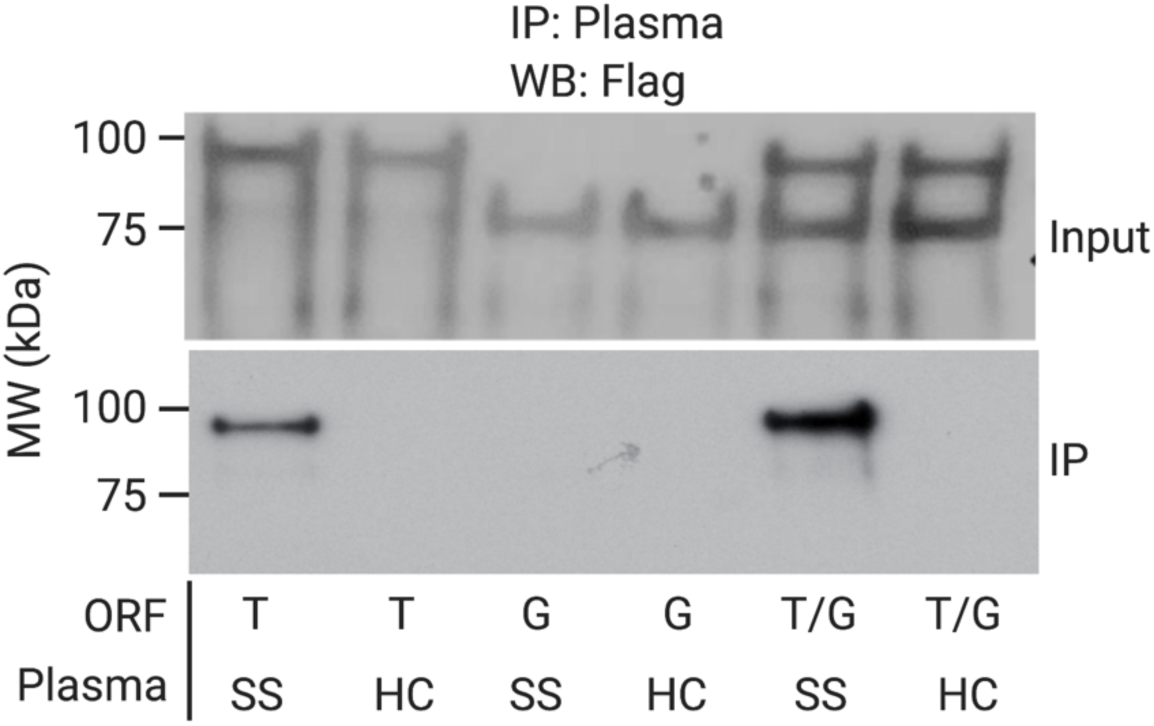
Two-plex translation and IP of TRIM21 and GAPDH. TRIM21 (T) and GAPDH (G) IVT-RNA-cDNA were translated either separately or together and then subjected to IP with healthy control (HC) or Sjogren’s Syndrome (SS) plasma. Analysis was by immunoblotting with the M2 antibody that recognizes the common FLAG epitope tag that links the HaloTag to the protein.

**Fig. S4.**
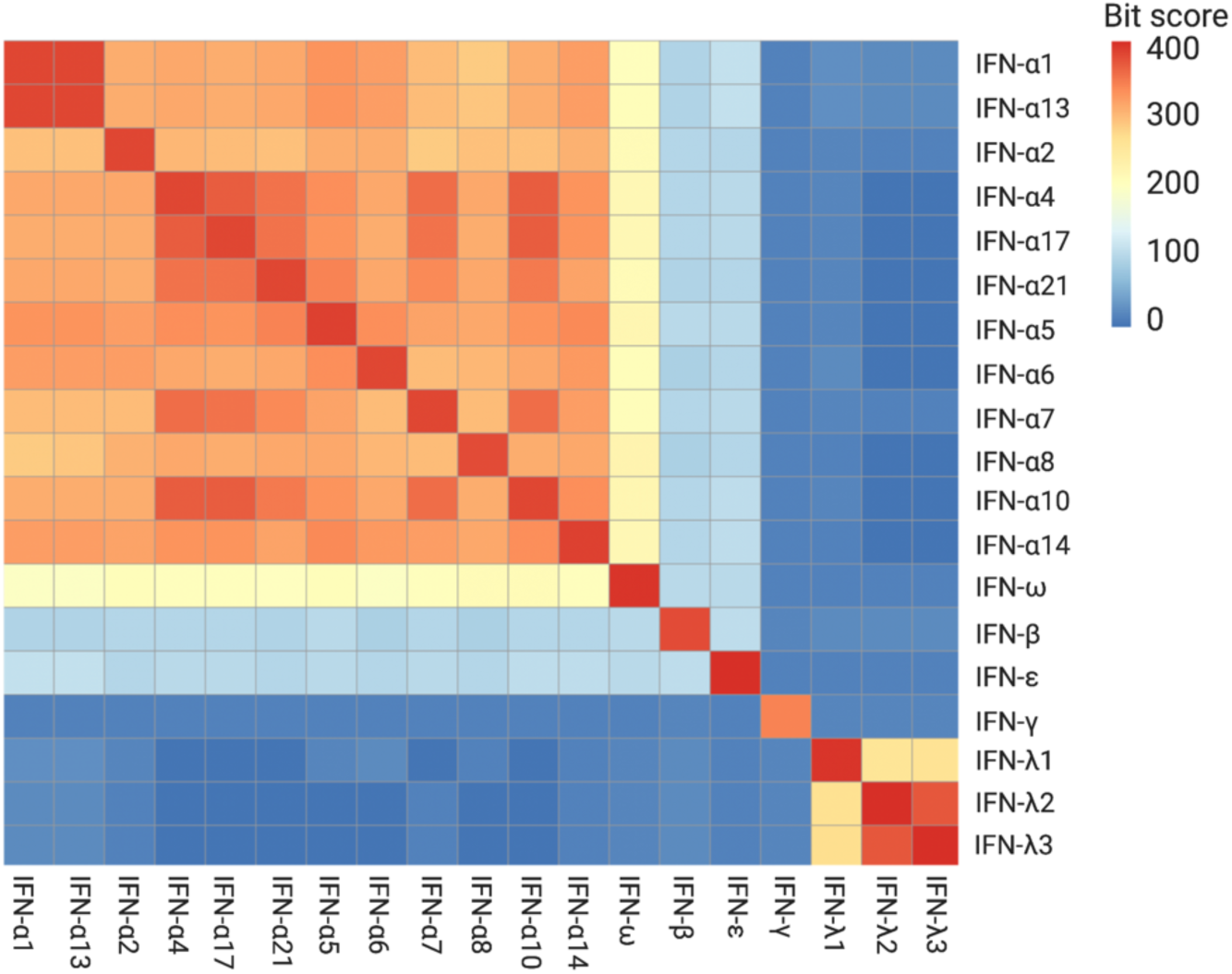
Sequence homology of interferons. Pairwise blastp alignment bitscore matrix for all interferon (IFN) proteins shown in Fig. 5D.

**Table S1.** Severe COVID-19 patients and control study participants. Individuals’ ages were provided to investigators as intervals to protect identities of study participants.

| Study Population |  |  |  |  |  |  |  |
| --- | --- | --- | --- | --- | --- | --- | --- |
| Group |  | # | Age | Sex | Black | White | Other |
| **Severe COVID-19** | Died | 17 | 67 (27,87) | F: 8, M: 9 | 9 | 4 | 4 |
|  | Ventilated | 13 | 67 (27,82) | F: 9, M: 4 | 4 | 4 | 5 |
|  | Got O2 | 22 | 52 (27,82) | F: 9, M: 13 | 8 | 8 | 6 |
|  | No O2 | 3 | 46 (22,49) | F: 0, M: 3 | 0 | 3 | 0 |
| **COVID-19 Controls** | Mild/Mod | 10 | 35 (19,55) | F: 6, M: 4 | 0 | 8 | 2 |
|  | Healthy Control | 10 | 41.5 (22,66) | F: 3, M: 7 | 3 | 5 | 2 |
| **Myositis** | Inclusion Body | 10 | 53.9 (43.6,60.6) | F: 7, M: 3 | 1 | 7 | 2 |
|  | Healthy Control | 10 | 36.5 (20,60) | F: 5, M: 5 | 2 | 8 | 0 |

**Table S2.**
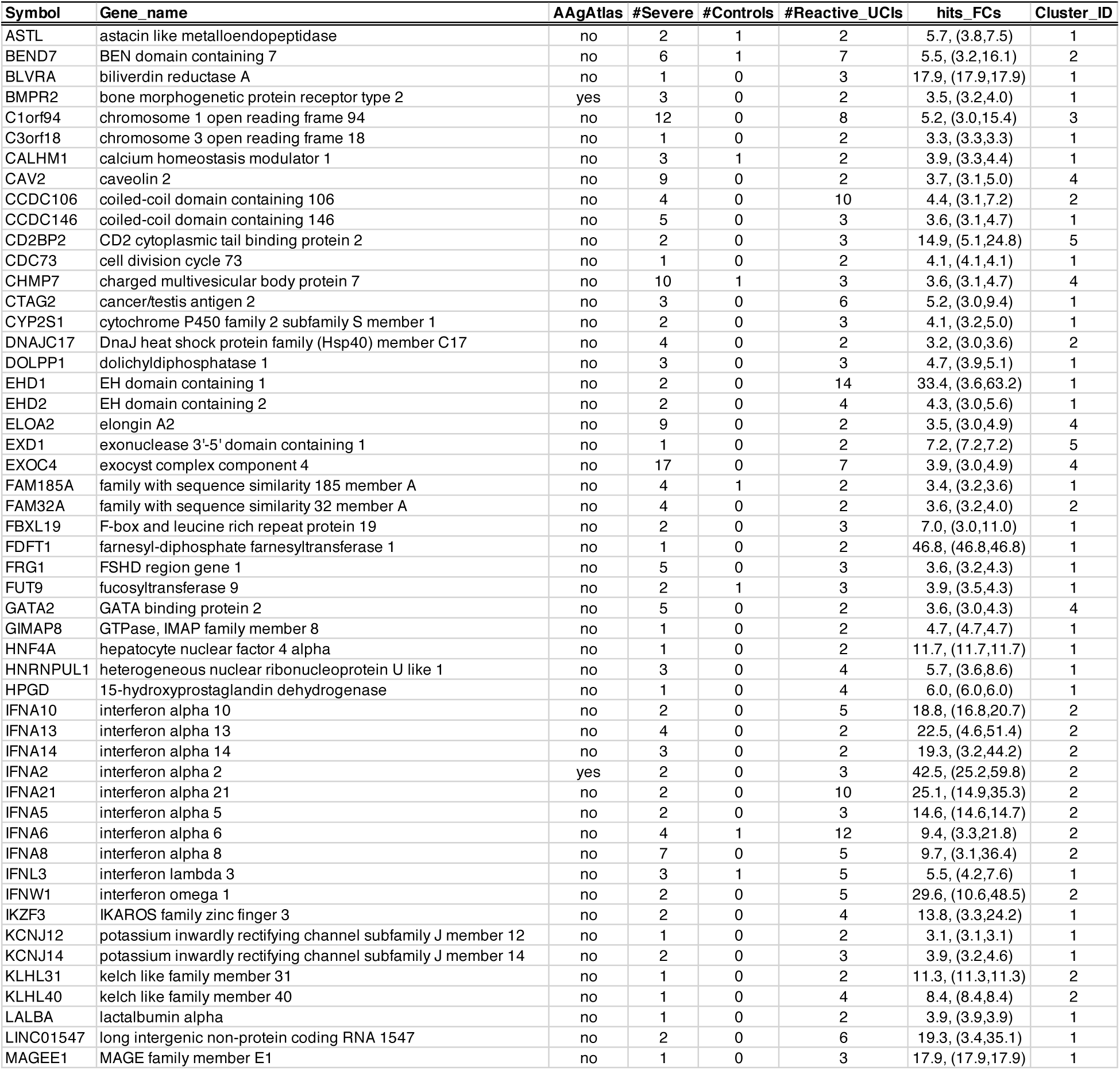
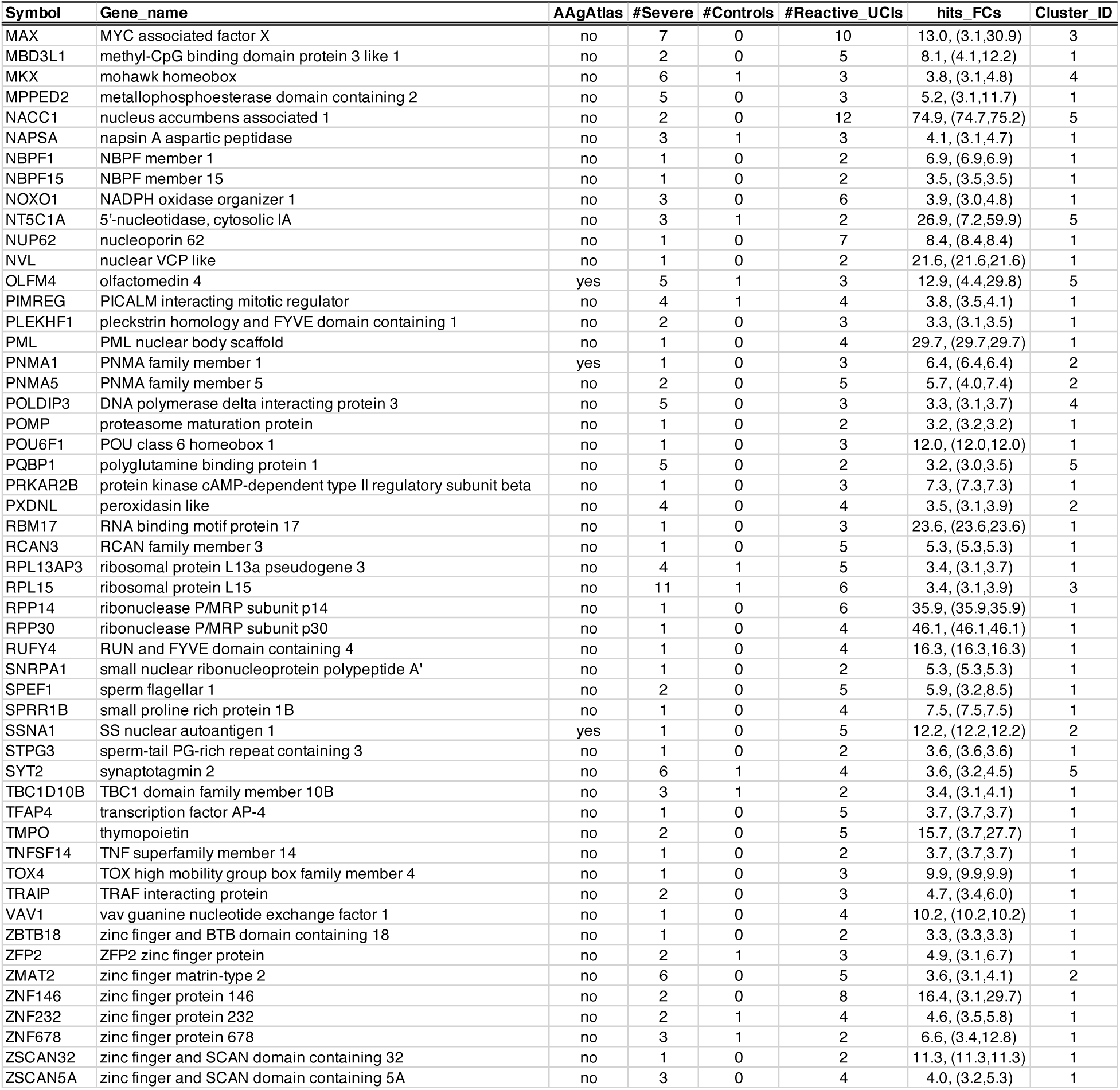
Proteins reactive in severe COVID-19 patients. Symbol, gene symbol. AAgAtlas, is protein listed in AAgAtlas 1.0? #Severe, number of severe COVID-19 patients with reactivity to at least one UCI. #Controls, number of control donors (healthy or mild-moderate COVID-19) with reactivity to at least one UCI. #Reactive_UCIs, number of reactive UCIs associated with given ORF. Hits_FCs, mean and range (minimum to maximum) of per-ORF maximum hits fold-change observed among the patients with the reactivity. Cluster_ID, row cluster defined by **Fig 4B**.

**Table S3.**
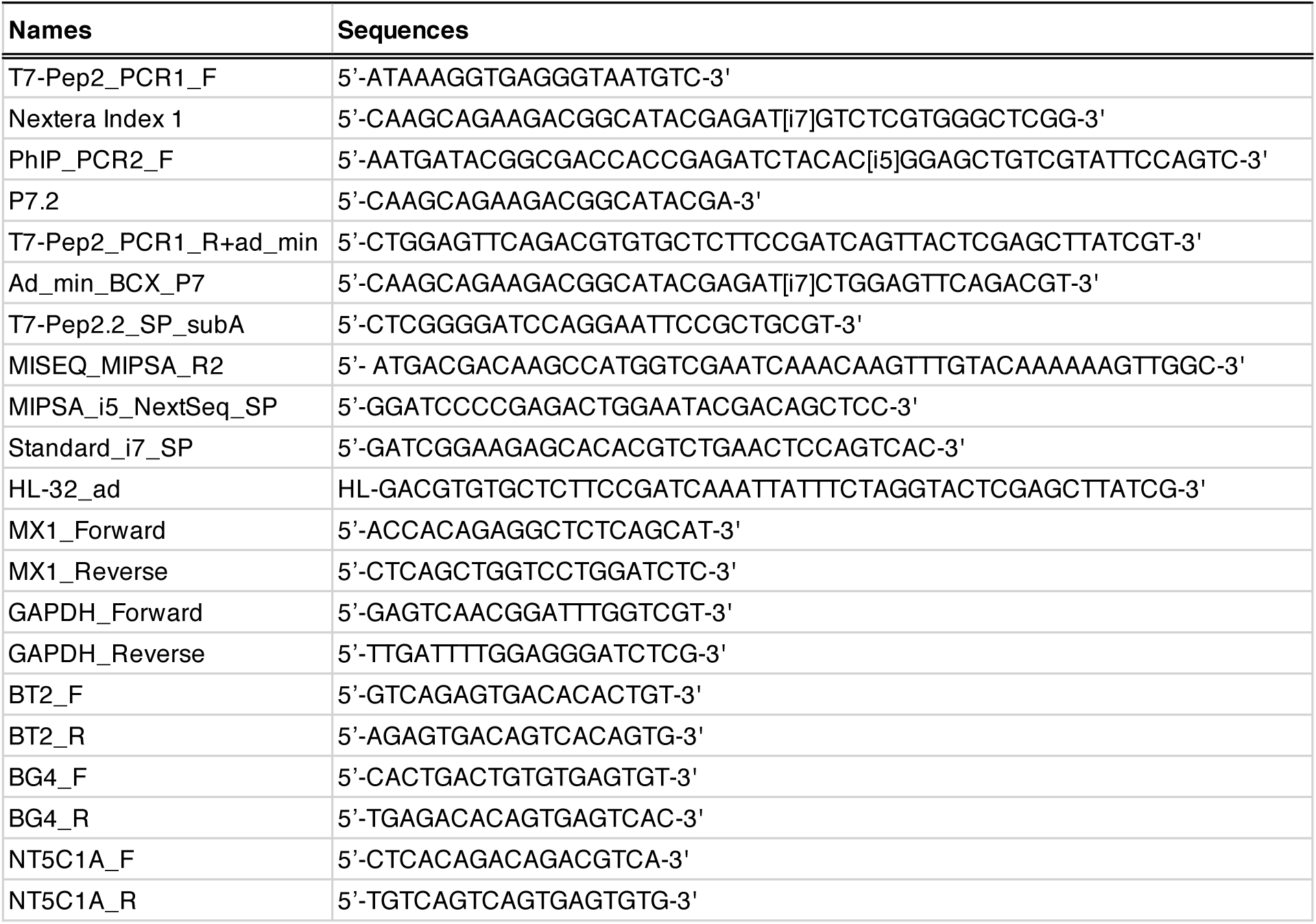
Primer sequences used in this study.

**Data S1. Hits fold-change MIPSA data matrix for UCIs of reactive proteins in severe COVID-19 patients**. (separate file)

